# From berries to brain: Assessing the impact of (poly)phenols in the MPTP mouse model of Parkinson’s disease

**DOI:** 10.1101/2025.08.16.670034

**Authors:** Rafael Carecho, Ana Filipa Raimundo, María Ángeles Ávila-Gálvez, Hugo Tainha Lopes, Catarina de Oliveira Sequeira, Carlos Pita, Catarina Pinto, Diogo Carregosa, Alicia Marín, Antonio González-Sarrías, Juan Carlos Espín, Sofia Azeredo Pereira, Andreia Neves Carvalho, Teresa F. Pais, Natasa Loncarevic-Vasiljkovic, Cláudia Nunes dos Santos

## Abstract

The growing burden of chronic neurodegenerative diseases (NDs), particularly Parkinson’s disease (PD), prompts the need for effective preventive strategies and treatments. Dietary (poly)phenols have emerged for their neuroprotective potential. This study investigated a (poly)phenol-enriched diet, comprising a berry mixture, to counteract key PD hallmarks in 1-methyl-4-phenyl-1,2,3,6-tetrahydropyridine (MPTP)-intoxicated mice and elucidate the phenolic metabolic fingerprint underlying these effects. The berry-enriched diet prevented motor deficits in the MPTP mice model, preserved dopaminergic neurons in the midbrain, reduced glial activation and gene expression of inflammatory cytokines. Notably, berries also attenuated macrophage infiltration observed in the substantia nigra 7 days post-MPTP. Metabolomic analysis revealed distinct phenolic signatures in plasma and brain tissue between standard- and berry-fed mice. Overall, this pioneering study provides compelling evidence that a (poly)phenol-enriched diet may play a protective role in neurodegenerative disorders, highlighting its potential for future strategies to prevent or slow PD progression.

## Introduction

The inevitable process of ageing brings with it a host of cognitive and physical challenges and a greater risk of developing neurodegenerative disorders (NDs), such as Parkinson’s disease (PD). The growing burden of PD across the globe, affecting more than 8.5 million people worldwide^1^, underscores the urgent need for enhanced preventive strategies and more effective treatments able to mitigate its devastating impact on PD patients’ lives.

The death of dopaminergic neurons in the substantia nigra pars compacta (SNpc) represents the main hallmark of PD. Although the underlying cellular mechanisms are still not fully understood, neuroinflammation and oxidative stress have been recognised as two of the main features of PD and key contributors to dopaminergic cell loss and, consequently, dopamine (DA) depletion in both familial and sporadic cases^2–4^. When exacerbated, the reactivity of microglia and astroglia, sustained by changes in their morphology and increased numbers, contributes to neuronal damage via the release of pro-inflammatory mediators and neurotoxic products^55^. In line with this, increased levels of pro-inflammatory cytokines have been detected in the dopaminergic regions and cerebrospinal fluid (CSF) of PD patients^6,7^, as well as in the brains of the animal models simulating the disease^8,9^. Furthermore, oxidative stress, characterised as an imbalance between the production of harmful free radicals and the ability to detoxify these reactive products, as those involving crucial components like glutathione (GSH) and cysteine (Cys)^10,11^, comprises another critical hallmark of PD.

Currently, PD remains incurable, with no effective therapies to delay or reverse the disease progression. Symptomatic therapeutic approaches currently used to restore DA levels and alleviate motor symptoms have been associated with long-term usage secondary effects^12,13^. Therefore, there is a pressing need for disease-modifying therapies and strategies to prevent or delay PD rather than strategies for symptomatic relief in later stages of the disease.

In this regard, diet is an emerging and attractive approach with increasing experimental support^14^. Dietary (poly)phenols, a big class of molecules abundant in fruits and vegetables, have emerged in the past years as promising compounds against PD since they are capable of mitigating both neuroinflammation and oxidative stress and improving cognitive and motor symptoms in different *in vivo* PD models (reviewed in ^15,16^).

Although the results obtained so far are promising, the research on (poly)phenol effects in PD lacks three important assets. The first one is that most of the studies so far employ (poly)phenols at high concentrations, relating more to the pharmacological approach (reviewed in ^15,16^). Considering that PD starts 10-20 years before the occurrence of the first symptoms^17^, this approach could have safety implications due to the high concentrations of compounds used for a prolonged period. Secondly, it is well described that ingested (poly)phenols pass through different metabolism steps, including phase I and II liver metabolism, and the metabolism by gut microbiota^18^, where the compounds that ultimately reach circulation and organs are predominantly low-molecular-weight (poly)phenol metabolites (LMWPMs) rather than the original parent compounds^19,20^. However, the data revealing the exact LMWPMs reaching the brain, which might be the effectors of the improvements observed in motor behaviour, neuroinflammation and oxidative stress in *in vivo* PD studies, is still missing. The third asset is that most of the studies performed so far employ either a single phenolic compound or, rarely, a plant extract of a single plant species^15,16,21^. Keeping in mind that each parent (poly)phenol compound can give rise to numerous LMWPMs^20^, using a dietary approach will comprise an entire palette of different (poly)phenols rather than a single parent compound, thus ensuring a multitude of LMWPMs reaching the circulation.

Therefore, an optimal strategy would encompass a preventive dietary approach incorporating a daily intake of diverse (poly)phenols. Such approach could offer the most advantageous outcomes, ensuring sustainability and safety. To outbalance this knowledge gap, we performed a study using a berry-enriched diet consisting of a mixture of blueberry, raspberry, and blackberry in a preventive approach for 6 weeks before inducing PD-like phenotype in mice, using 1-methyl-4-phenyl-1,2,3,6-tetrahydropyridine (MPTP) intoxication. The majority of the in vivo MPTP studies use either acute or sub-chronic injection protocols^22^. In our study, we opted for the acute MPTP paradigm as it has been shown to induce motor impairment far more than the sub-chronic one^22^. Berries were chosen as a well-documented source of (poly)phenols that can be seamlessly and safely integrated into a daily diet^24–26^. Our results indicate that berry supplementation can significantly improve MPTP-induced motor deficits, decrease dopaminergic cell death, and alleviate micro- and astrogliosis. Moreover, metabolomic analysis disclosed that the main LMWPMs occur in circulation and inside the brain. Our research provides a step further, a thorough understanding of how dietary (poly)phenols can proactively influence the primary pathological processes in the brain affected by PD, thereby contributing to preventing or delaying the onset of PD symptoms.

## Results

### A berry-enriched diet counteracts MPTP-induced corticosterone increase in urine without affecting food intake or weight gain in mice

Mice were fed either a standard or a berry-enriched diet for six weeks prior to MPTP administration (**Figure 1A**). The proximate compositions of standard and berry-enriched diets were determined (**Table S1, Supplementary Information**). The dominant components of both diets consisted of carbohydrates, water, and proteins, with no statistically significant differences between samples. Supplementation with the berry mixture did not significantly alter the nutritional value of the diet. The standard diet is a plant-based diet mainly composed of ground wheat and corn, wheat middlings, and soybean meal supplemented with minerals and vitamins. The manufacturers’ characterisation describes the inclusion of soybean meal resulting in an expected isoflavone range of 225-340 mg/kg diet (daidzein + genistein aglycone equivalents) that corresponds, on average, to a daily consumption of 0.9 - 1.36 mg (daidzein + genistein aglycone equivalents). To confirm that the consumption of an 8% berry-enriched diet increased the daily uptake of (poly)phenols, we characterised the phenolic profile of both diets and analysed the plasma metabolome of the animals. Total phenolic content (TPC) analysis demonstrated a significantly higher content of phenolic compounds in the 8% berry-supplemented diet (471± 186 mg GAE/100 g) in comparison with the standard diet (184 ± 47.6 mg GAE/100 g) (**Figure S2, Supplementary Information**). In the standard diet, we detected di-*O*-caffeoylquinic acid and caffeic acid, while a more complex mixture of phenolic compounds was present in the berry-enriched diet (**Tables S2 and S3, Supplementary Information**). During the initial 6 weeks of the intervention, no differences were noted in food intake or weight gain between groups (**Figures 1B and 1C**).

**Figure 1.**
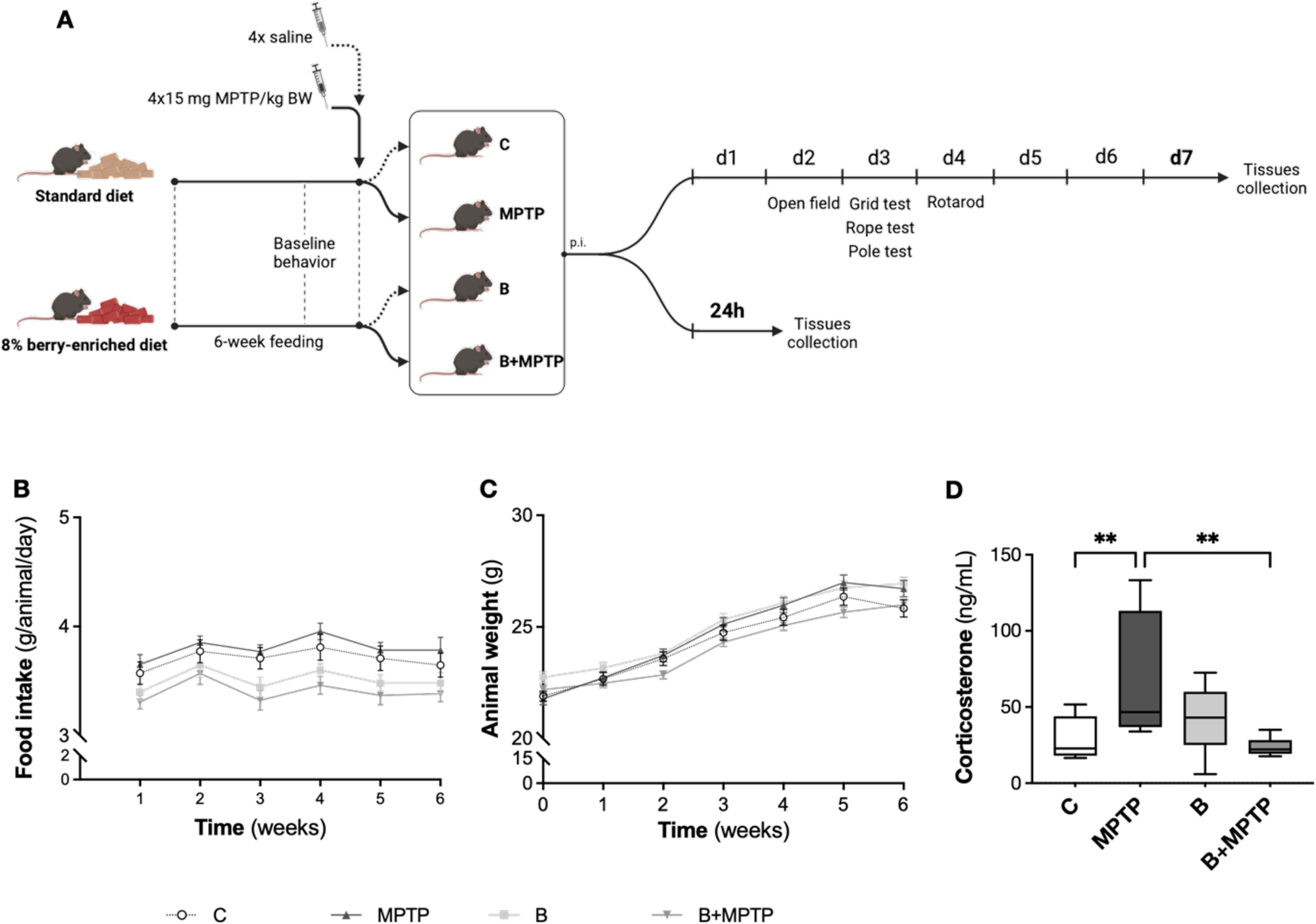
Effect of the berry-enriched diet on food intake, animal weight, and corticosterone levels. (**A**) Schematic diagram illustrating the experimental design and timeline of the study. Male C57BL/6J mice were fed for 6 weeks, either with standard chow or an 8% berry-enriched diet. Following this period, half of the mice of each diet group were submitted to four intraperitoneal-injections (IP) of 15 mg of MPTP/kg of body weight (black arrow) to induce a Parkinson’s disease-like phenotype (MPTP group and B+MPTP group), while the rest received four saline injections (dash arrow) (C group and B group). Two groups of animals were sacrificed at two time points, 24 hours or 7 days after the injections. Baseline behaviour was conducted before the injections, and behaviour assessments were performed between day 2 and day 4 post-injections, as shown. Daily food intake (**B**) and body weight (**C**) for the initial 6 weeks of the intervention. (**D**) Urine corticosterone levels 5 days post-MPTP injection. Statistical significance: **p<0.01. C – Control group; MPTP – MPTP group; B – Berry group; B+MPTP – Berry+MPTP group; d – day; p.i. – post-injection.

Mice of all four groups were sacrificed either 24h or 7d post-injections (**Figure 1A**). Importantly, a test evaluating the potential interference of a berry diet (and its molecules) on MPTP metabolism was previously performed, due to evidence that some (poly)phenols can inhibit monoamine oxidase B (MAO-B)^23,24^, the enzyme responsible for converting MPTP to MPP^+^, the active toxic compound. As such, the blood levels of MPTP and MPP^+^ in the striatum were quantified. No differences between mice fed a standard diet and mice fed a berry-enriched diet were detected regarding MPTP or MPP^+^, implying the absence of relevant interference of berry feeding with the metabolism of MPTP (**Figure S1, Supplementary Information**).

To evaluate stress and metabolic response, the levels of corticosterone were evaluated in the mice urine (**Figure 1D**). Notably, our findings suggest that MPTP induces a physiological stress response in mice, as evidenced by the significantly increased corticosterone levels in MPTP-administered mice compared to the control levels, while a berry-enriched diet completely abolishes this response (**Figure 1D**).

### Mouse brain and plasma phenolic metabolome are changed upon a berry-enriched diet

Since it is known that the most abundant (poly)phenols found in circulation upon flavonoid and chlorogenic acid intake are the LMWPMs^20^, we focused our target metabolomic approach on 132 (poly)phenols and LMWPMs (**Table S4, Supplementary Information**). Of those, 40 were detected, either in the plasma (39) and/or in the perfused brains (17) of C and B groups (**Table 1**). Interestingly, phloroglucinol was not detected in plasma, but was detected in the brains of both groups (**Table 1**). Resorcinol, benzoic acid, hydroxybenzoic acid, dihydroxybenzoic acid, phloroglucinaldehyde, hippuric acid, caffeic acid, 3-(4′-hydroxyphenyl)propanoic acid, ferulic acid, caffeic acid sulfate, dihydrocaffeic acid, and vanillic acid sulfate were detected and quantified in the plasma, yet no differences were found between groups (**Figure S3, Supplementary Information**). Meanwhile, some metabolites were exclusively detected in circulation in berry-fed animals, such as 5-(3’,4’-dihydroxyphenyl)-*γ*-valerolactone sulfate, 5-(phenyl)-*γ*-valerolactone sulfate, and methyl-*O*-epicatechin-sulfate (**Table 1**), a clear indication of flavan-3-ol intake that corroborates the high variety of flavonoids present in the berry-enriched diet.

**Table 1.**
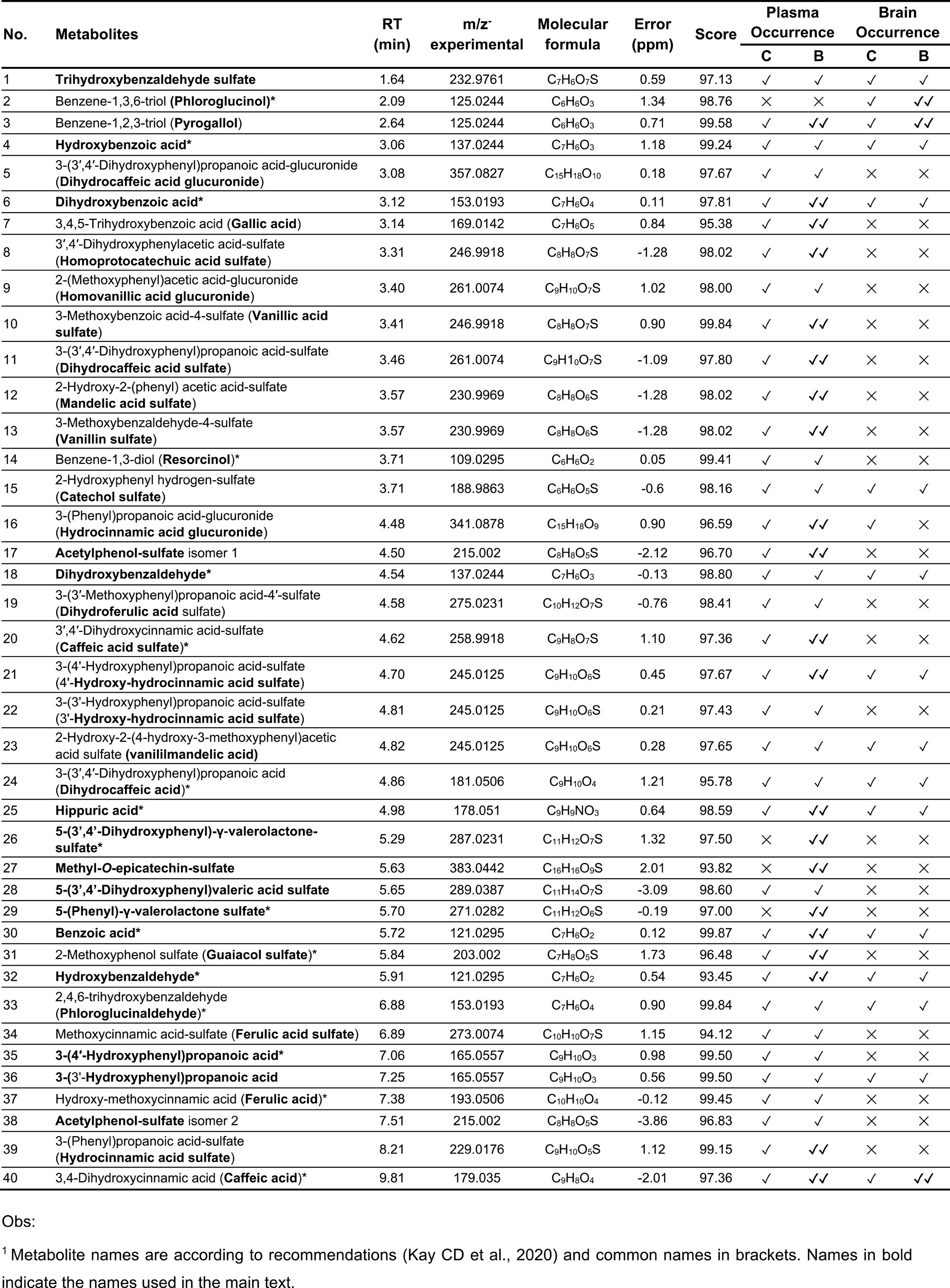
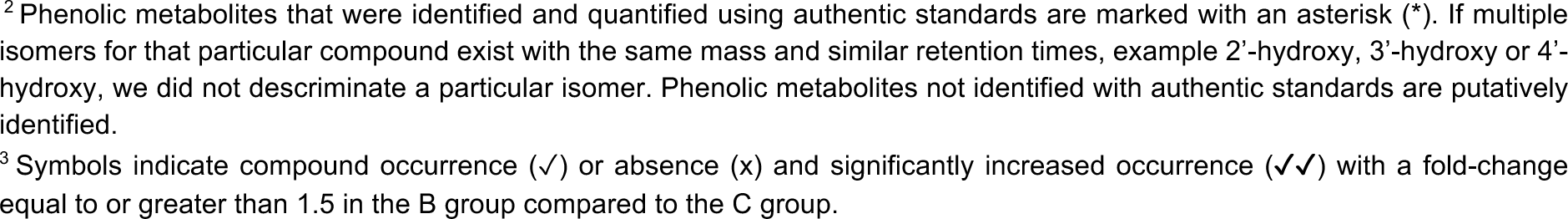
(Poly)phenols and derived metabolites in plasma and/or perfused brain of Control (C) and Berry (B) groups following 7 weeks of feeding^1,2,3^.

The phenolic metabolomic profile of plasma and perfused brain samples after 7 weeks of berry feeding resulted in an increase in the overall concentration of metabolites circulating in the B group animals compared with the C group. Relative quantification of the peak areas of each metabolite showed a fold-change (FC) increase of 1.5 or more, on 21 out of 39 metabolites detected in the plasma of the berry-fed animals compared to the standard diet (**Table 1** and **Figure 2A**). The plasma PCA revealed a distinct clustering pattern between the two diets, primarily driven by the distinct occurrence of metabolites (**Figure 2B**). Despite an overlapping region, the PCA plot separation suggests that the overall metabolomic profiles of these groups were different, providing the animals with a distinct starting point before being challenged by MPTP intoxication. Among the 21 metabolites (≥1.5-fold-change B/C), quantification was performed for those with available standards. Quantification confirmed increased circulating concentrations in the B group compared with the C group for the metabolites 5-(3’,4’-dihydroxyphenyl)-γ-valerolactone sulfate (**Figure 2C**), 5-(phenyl)-*γ*-valerolactone sulfate (**Figure 2D**), guaiacol sulfate (**Figure 2E**) and pyrogallol (**Figure 2F**).

**Figure 2.**
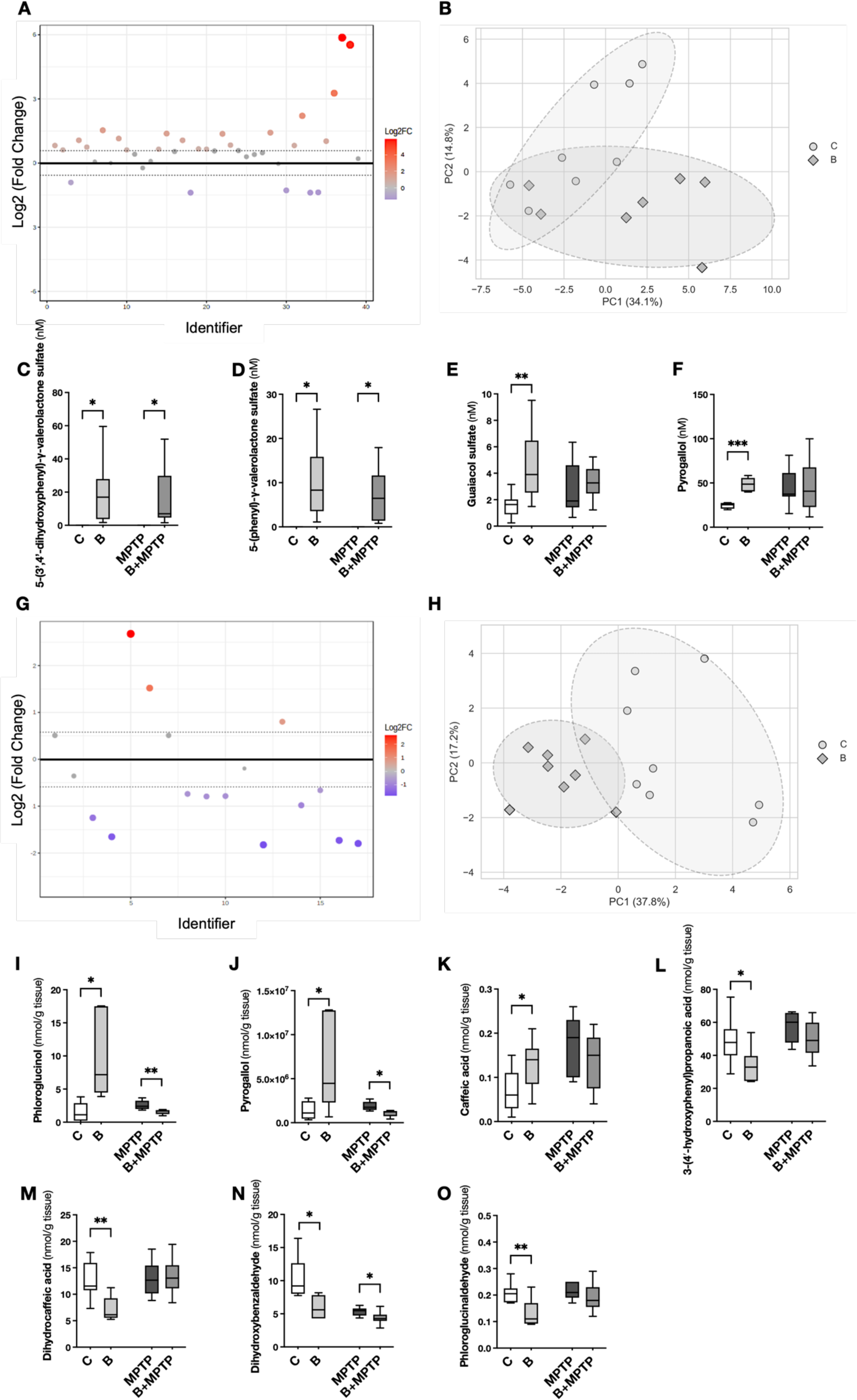
Univariate and multivariate analysis of phenolic metabolites in plasma and perfused brain. (**A**) Fold-change analysis presented as log2(fold-change B/C), highlighting the metabolites higher than log2(1.5-fold-change B/C) detected in the plasma. (**B**) Two-dimensional PCA score plot of both B and C groups considering the metabolites detected in circulation after 7 weeks of feeding. Quantification of (**C**) 5-(3’,4’-dihydroxyphenyl)-γ-valerolactone sulfate, (**D**) 5-(phenyl)-γ-valerolactone sulfate, (**E**) Guaiacol sulfate and (**F**) Pyrogallol in the plasma. (**G**) Fold-change differences between the berry-enriched diet and control diet in the brain. (**H**) Two-dimensional PCA score plots of both B and C groups considering the metabolites detected in the brain after 7 weeks of feeding. Quantification of (**I**) Phloroglucinol, (**J**) Pyrogallol, (**K**) Caffeic acid, (**L**) 3-(4′-hydroxyphenyl)propanoic acid, (**M**) Dihydrocaffeic acid, (**N**) Dihydroxybenzaldehyde, and (**O**) Phloroglucinaldehyde in the brain. *n*=8-10. Statistical significance: *p<0.05, **p<0.01, ***p<0.001. C – Control group; MPTP – MPTP group; B – Berry group; B+MPTP – Berry+MPTP group.

The analysis of metabolites detected in the brain of both C and B groups revealed insightful trends. Out of 17 metabolites identified in the brain, only phloroglucinol, pyrogallol, and caffeic acid exhibited a 1.5-fold increase in B animals compared to C animals. In contrast, dihydroxybenzoic acid, catechol sulfate, dihydroxybenzaldehyde, phloroglucinaldehyde, trihydroxybenzaldheyde sulfate, 2-hydroxy-2-(4-hydroxy-3-methoxyphenyl)acetic acid sulfate, dihydrocaffeic acid, 3-(4′-hydroxyphenyl)propanoic acid and hydroxycinnamic acid sulfate were decreased (FC<1.5) in berry feed animals (**Figure 2G**). PCA (**Figure 2H**) maximised the overall distribution and clustering of the metabolomic data from the C and B groups. Interestingly, considering all four groups, we observed a statistically significant increase in the concentrations of phloroglucinol (**Figure 2I**), pyrogallol (**Figure 2J**), and caffeic acid (**Figure 2K**) in the B group compared to the control. On the contrary, 3-(4′-hydroxyphenyl)propanoic acid (**Figure 2L**), dihydrocaffeic acid (**Figure 2M**), dihydroxybenzaldehyde (**Figure 2N**) and phloroglucinaldehyde (**Figure 2O**) levels decreased in the B group compared to control animals. Differences were found between MPTP and B+MPTP groups, on phloroglucinol (**Figure 2I**), pyrogallol (**Figure 2J**), and dihydroxybenzaldehyde (**Figure 2N**). No differences were denoted for benzoic acid, hydroxybenzoic acid, dihydroxybenzoic acid, hydroxybenzaldehyde or hippuric acid (**Figure S4, Supplementary Information**). Altogether, this shift in the brain’s metabolomic profile, when considered alongside the findings in plasma metabolites, points out the impact of the berry-rich diet on circulating metabolites and brain metabolome.

### A berry-enriched diet attenuates motor dysfunction in MPTP-intoxicated mice

Motor disabilities are one of the most significant symptoms of PD. We observed that MPTP intoxication caused severe motor impairments as evaluated by behavioural tests as early as 48h post-injection. Mice were submitted to the open field test to assess spontaneous and voluntary movement. Both track and occupancy plots (**Figure 3A**) disclosed a robust contrast regarding overall motor activity between control and MPTP-treated mice. Notably, the overall activity time was significantly lower in MPTP mice, while berry supplementation could not counteract this effect (**Figure 3B**). Moreover, although not significant, MPTP-treated mice seemed to spend less time in the outer field of the arena compared to C and B+MPTP groups (**Figure 3C**). The difference in the ambulatory time was reflected in the total travelled distance (**Figure 3D**) and the number of centre crossings (**Figure 3E**). MPTP-treated mice were particularly slower than the C group, and interestingly, B+MPTP presented the lowest mean velocity (**Figure 3F**). The open field test also allows for addressing exploratory behaviour and as a measure of anxiety, besides the increased latency to leave the centre (**Figure 3G**). Upon object introduction, B+MPTP mice displayed the highest latency in approaching the object (**Figure 3H**) and the lowest number of object approaches (**Figure 3I**). No differences between the groups were noted in the object exploration time (**Figure 3J**). Noteworthy, when evaluating a model with motor impairments, any consideration related to the anxiety levels of mice should be interpreted with caution since a decrease in exploratory activity and fewer centre crossings might be due to overall reduced motor behaviour caused by MPTP intoxication and not to increased anxiety levels.

**Figure 3.**
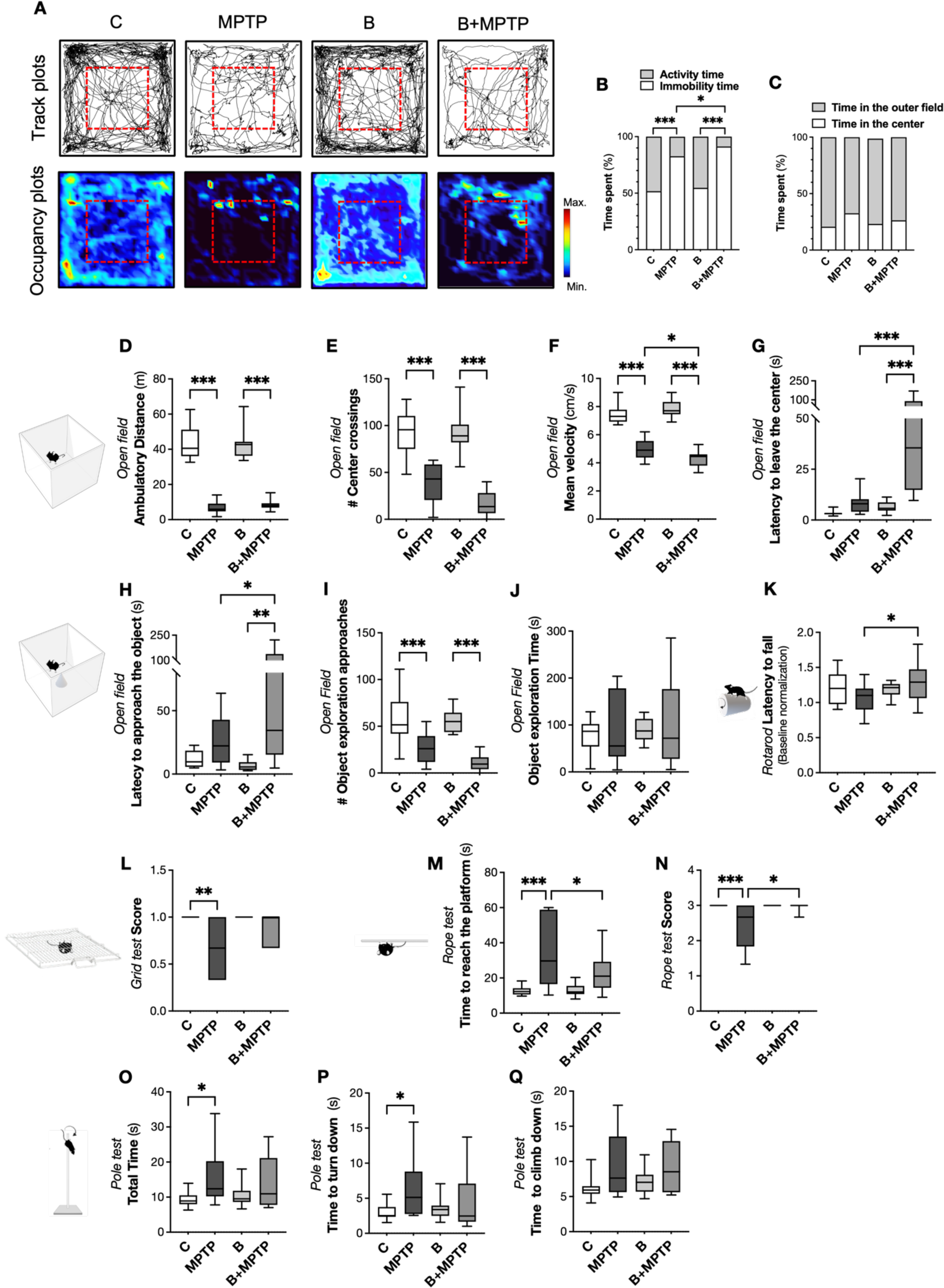
Behavioural testing of motor function between days 2 and 4 post-MPTP injections. (**A**) Representative track plots and heat maps with occupancy plots of mice in the open arena. (**B**) Activity vs immobility time (%) and (**C**) time spent in the outer field vs. the centre (%). (**D**) Ambulatory distance, (**E**) number of centre crossings, (**F**) mean velocity, and (**G**) latency to leave the centre. Following 20 min in the open field, a novel object was introduced, in the centre of the arena, for 10 min. (**H**) Latency to approach the object, (**I**) number of object exploration approaches, and (**J**) object exploration time. (**K**) Rotarod acceleration protocol (baseline normalisation). Grid test (**L**) score. Rope test parameters: (**M**) time to reach the platform, and (**N**) score. Pole test parameters: (**O**) total time, (**P**) time to turn, and (**Q**) time to climb down. *n*=12-16 per group. Statistical significance: *p<0.05, **p<0.01, ***p<0.001. C – Control group; MPTP – MPTP group; B – Berry group; B+MPTP – Berry+MPTP group

Fine motor tests evaluating motor coordination and strength were also performed. Remarkably, in the accelerating-speed rotarod protocol, the time on the rod was significantly lengthened by berry feeding in MPTP-treated mice, whose performance was even better than control groups (**Figure 3K**). In the grid test, MPTP mice showed a significant increase in fall incidence (meaning reduced score), which was improved in a minor, non-significant manner in the B-MPTP group (**Figure 3L**).

In the rope test, animals in the MPTP group exhibited increased latency to reach the platform relative to control animals (**Figure 3M**), and obtained significantly lower scores, indicative of reduced task completion frequency (**Figure 3N**). Notably, B+MPTP animals demonstrated motor performance more closely aligned with that of their respective controls, suggesting improved motor function in the execution of the task (**Figure 3M, N**).

Motor impairments in the MPTP group were also evident in the pole test, marked by a significant increase in the total time required to complete the task (**Figure 3O**). In detail, MPTP-treated mice took more time to turn downwards the vertical pole (**Figure 3P**) and to climb down (**Figure 3Q**). In this test, no protective effects were observed on the berry diet.

Behavioural deficits observed in MPTP-treated animals were consistent with the PD model, whereas a berry-enriched diet promoting the generation of LMWPMs prevented these symptoms, highlighting its potential to counteract PD-related motor impairments.

### A berry-enriched diet delays MPTP-induced dopaminergic neuronal damage in the midbrain

Acute administration of MPTP is known to induce dopaminergic neurodegeneration. Therefore, we initially evaluated cell death in the midbrain, focusing on the SNpc, 24h after MPTP exposure. In both MPTP and B+MPTP groups, cells labelled with Fluoro-Jade C, a marker of neuronal degeneration, were detected (**Figure 4A**). Notably, although not significant, the B+MPTP group tended to exhibit fewer positively labelled cells than the MPTP group (**Figure 4A**). The tendency observed in Fluoro-Jade C was reflected in TH protein levels in the midbrain following 24h post-injections, which decreased in the MPTP group (**Figure 4B**). Interestingly, the berry-enriched diet sustained MPTP-induced TH reduction, presenting levels close to those observed in the C group in the midbrain (**Figure 4B**). Regarding apoptotic cell death, no TUNEL-positive cells were detected either in MPTP or in the B+MPTP group (**Figure S5, Supplementary Information**), indicating that 24h post-MPTP intoxication, cells were not dying by apoptosis. Also, no alterations were observed in the midbrain levels of anti- and pro-apoptotic proteins (B-cell lymphoma-2 (Bcl-2) and Bcl-2-associated X (Bax, respectively) (**Figure 4B**). Additionally, no differences were observed in TH levels in the striatum (**Figure 4C**). These results suggest that berries might delay the severity of the MPTP phenotype by sustaining the TH levels in the midbrain, as the primary region affected in PD. In contrast, the striatum was not affected 24h after post-MPTP injection.

**Figure 4.**
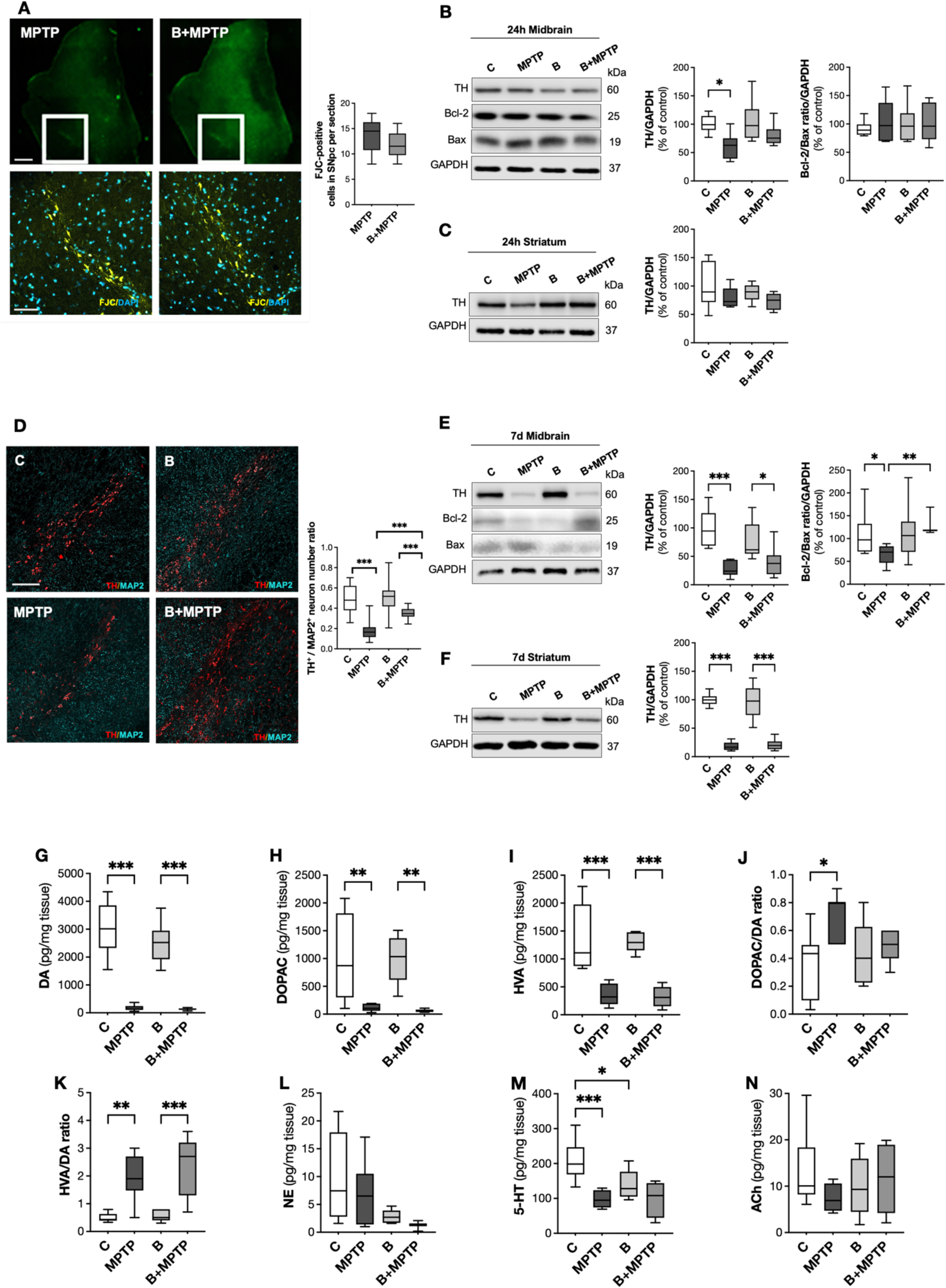
Evaluation of neurodegeneration and dopaminergic markers in both midbrain and striatum. (**A**) Fluoro Jade-C (FJC) assay of the midbrain (upper images of the panel). Lower images represent the zoomed sections of FJC-labelled degenerating neurons in the SNpc, with nuclei stained with DAPI (blue), 24h after the MPTP injections (*n*=4 per group). Average number of FJC-labelled cells in the SNpc per section. (**B**) Tyrosine Hydroxylase (TH), Bcl-2 and Bax protein levels in the midbrain at 24h post-injections (*n*=6-8 per group). (**C**) TH protein levels in the striatum at 24h post-injections (*n*=6-8 per group). (**D**) Representative images of TH/MAP2 double immunostaining of the SNpc 7d after the MPTP injections (*n*=4 per group). Average number of TH-positive neurons in the SNpc per section, normalised to the number of MAP2-positive neurons in the SNpc in the same section. (**E**) TH, Bcl-2 and Bax protein levels in the midbrain at 7d post-injections (*n*=6-8 per group). (**F**) TH protein levels in the striatum at 7d post-injections (*n*=6-8 per group). Striatal levels of (**G**) dopamine (DA), and its main metabolites (**H**) 3,4-dihydroxyphenylacetic acid (DOPAC) and (**I**) homovanilic acid (HVA), (**J**) DOPAC/DA and (**K**) HVA/DA ratios and the neurotransmitters (**L**) norepinephrine (NE), (**M**) serotonin (5-HT) and (**N**) acetylcholine (ACh), at 7d after MPTP administration (*n*=6-8 per group). Panel A upper images, scale bar: 500 µm; lower images, scale bar: 100 µm; Panel F, scale bar: 250 µm. Statistical significance: *p<0.05, **p<0.01, ***p<0.001. C – Control group; MPTP – MPTP group; B – Berry group; B+MPTP – Berry+MPTP group.

We next analysed the number of TH-positive cells in the SNpc 7d after MPTP injections to determine whether the berry-enriched diet can prevent dopaminergic neuronal death, considering that the majority of DA neuronal cell death occurs in the first 5-6 days after this regimen of acute MPTP intoxication^25^ (**Figure 4D**). We found that MPTP significantly reduced the number of TH-positive neurons in the SNpc of both MPTP and B+MPTP groups compared to the respective controls (C and B) (**Figure 4D**). Interestingly, berry supplementation significantly lessened the death of TH-positive cells in the SNpc of B+MPTP animals compared to the MPTP group (**Figure 4D**). This result was not reflected in the total TH protein level measured by western blot on the midbrain and striatum (**Figures 4E and 4F**). However, the berry diet attenuated the MPTP-induced Bcl-2/Bax ratio change, implying a positive effect of berries on hampering apoptosis (**Figure 4E**).

Since MPTP induced dopaminergic neuron loss, disruption in DA-related pathways was expected. We observed that MPTP induced a clear reduction in the levels of catecholamines such as DA, 3,4-dihydroxyphenylacetic acid (DOPAC), homovanillic acid (HVA) in the striatum (**Figures 4G, 4H, 4I**). Nonetheless, no significant difference was seen between MPTP and B+MPTP groups. Interestingly, MPTP intoxication markedly increased the ratios of DOPAC/DA, suggesting an increased DA turnover in the MPTP group, while the B+MPTP group showed no differences compared with the respective controls (**Figure 4J**). The HVA/DA ratio increased in the two groups of MPTP-treated animals (**Figure 4K**). Both DOPAC and HVA, besides being endogenously produced as DA metabolites, are also metabolites resulting from the gut microbiota (poly)phenols catabolism^20^. However, berry feeding did not affect their brain levels and, thus, could not compensate for the MPTP-induced disruption of catecholamine synthesis. No significant differences for DA, its metabolites and norepinephrine (NE) were observed between the C group and B group (**Figures 4G, 4H, 4I, and 4L**), suggesting that the berry-enriched diet did not affect catecholamine levels in normal mice. However, striatal serotonin (5-HT, 5-hidroxitriptamina) levels significantly decreased in the MPTP and berry-fed groups (**Figure 4M**). No statistical differences in acetylcholine (ACh) levels were noted in any experimental group (**Figure 4N**).

### A berry-enriched diet influences cysteine-related thiolome

The cellular low molecular thiols network (thiolome) is described to interfere in PD pathogenesis since it plays a crucial role in protecting neurons during oxidative stress^26^. GSH has a vital role in the thiolome, acting as a robust antioxidant defence system, maintaining redox homeostasis, and preventing oxidative damage to cellular components. GSH is synthesised in the cytosol from its constituents Cys, glutamate, and glycine (Gly)^27^. A PCA was performed considering the levels of the different fractions quantified in the brain of Cys, GSH and CysGly, a breakdown product of GSH catabolism, and an important source of Cys)^27^. A clear separation between C and MPTP groups was observed in the acute response to MPTP (24h timepoint) (**Figure 5A**). A distinct thiolome profile was also observed when comparing the C and B groups. The B+MPTP group could not be differentiated from the MPTP group, although B+MPTP appeared more dispersed. Importantly, the B+MPTP cluster overlapped the B group (**Figure 5A**). At 7d, there was no clear separation between the groups (**Figure 5B**). As reflected in the PCA profile, the late response (7 days post-MPTP) showed significant changes in CysGly dynamics, while the overall group profiles remained largely comparable (**Figure S6A-I, Supplementary Information)**.

**Figure 5.**
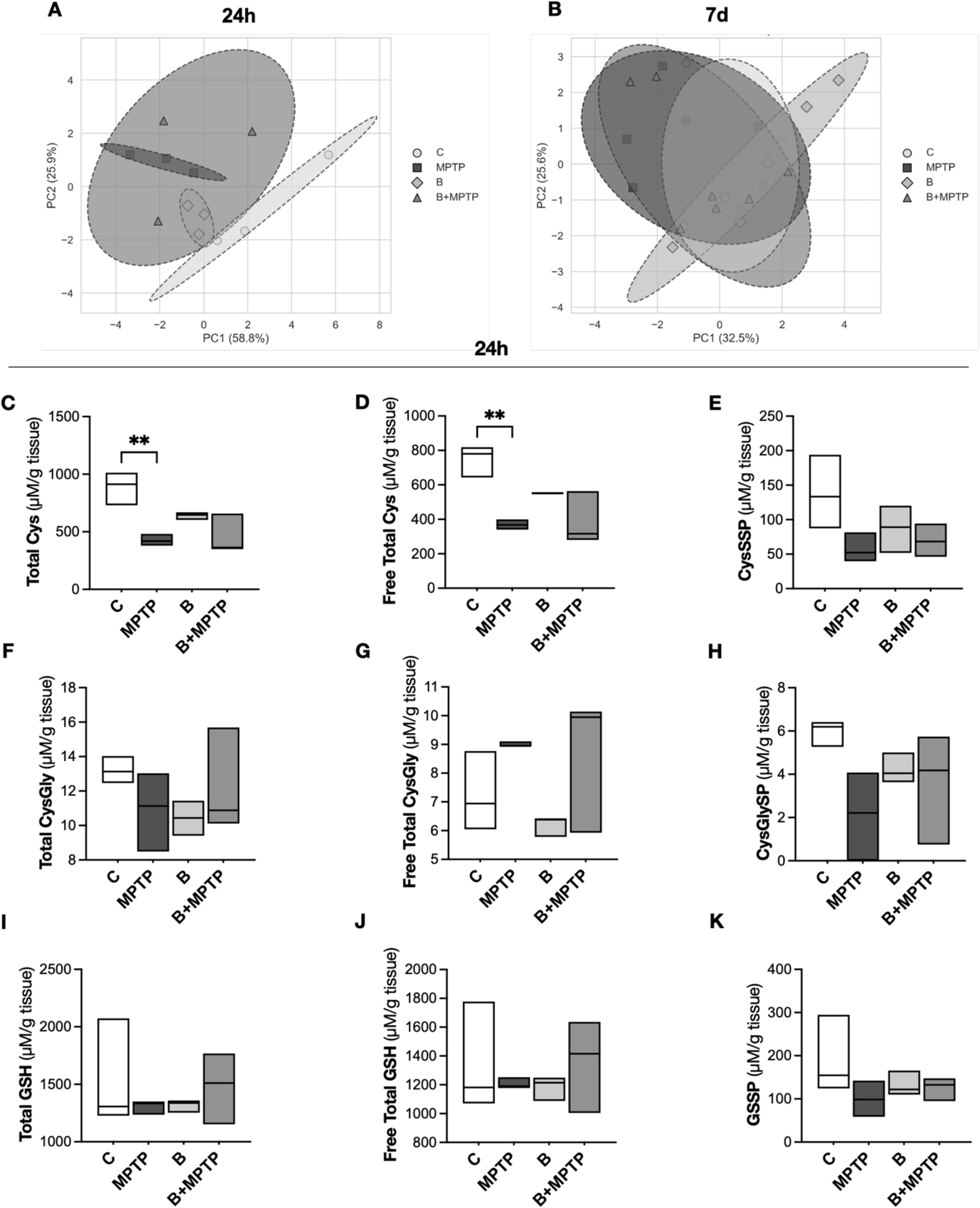
Cysteine-related thiols in the brain 24h post-MPTP injections. Principal Component Analysis (PCA) with all different fractions of cysteine (Cys), glutathione (GSH) and cysteineglycine (CysGly) quantified in the brain at (**A**) 24h and (**B**) 7d. Thiolome profile at 24h: (**C**) Total Cys; (**D**) Free total Cys; (**E**) Cysteinylated proteins (CysSSP); (**F**) Total CysGly; (**G**) Free total CysGly; (**H**) Cysteinylglycinylated proteins (CysGlySP); (**I**) Total GSH; (**J**) Free total GSH; (**K**) Glutathionylated proteins (GSSP). *n*=3 per group. Statistical significance: **p<0.01: comparison with C group. C – Control group; MPTP – MPTP group; B – Berry group; B+MPTP – Berry+MPTP group.

In turn, the analysis of the thiolome profile in the mouse brain 24h post-MPTP injections demonstrated that MPTP interfered mostly with the total levels of Cys by decreasing both free total Cys and protein-bound (CysSSP) fractions (**Figures 5C, 5D, and 5E**). No significant differences in CysGly or GSH levels were observed between groups at 24h across the three fractions: total, free total, and glutathionylated proteins (GSSP) (**Figures 5F–K**). Considering the subtle variation in clustering at 24h, we assessed several genes implicated in the cellular defence system against oxidative stress and toxicity in the brain, including *Gclm*, involved in GSH biosynthesis; *Gclc*, crucial for the first step of GSH synthesis; *Nfe2l2* (Nrf2), a key regulator of antioxidant defence mechanisms; and *Nqo1*, involved in detoxification of potentially harmful quinones. Unexpectedly, as part of a compensatory antioxidant response to counteract oxidative stress, helping to mitigate MPTP neurotoxic effects, no significant differences were observed in the mRNA expression of *Gclm*, *Gclc*, *Nfe2l2* (*Nrf2*), and *Nqo1* in any of the brain structures analysed (**Figure S6J-M, Supplementary Information)**.

### Microglial cell reactivity induced by MPTP is ameliorated in MPTP-treated mice fed with berries

Chronic microglial activation contributes to neurodegeneration, further propelling the pathogenesis of PD. We confirmed a marked increase in the number of Iba1-positive cells, in the SNpc, striatum, and motor cortex 24h after the MPTP injections, as well as an increase in microglia soma size (**Figures 6A-6C**). Impressively, the berry-enriched diet prevented this MPTP-induced microglia reactivity in all three analysed brain structures regarding cell number and their soma size (**Figures 6A-6C**). Interestingly, berry-feeding *per se* promoted a slight but significant increase in microglia soma size in all three brain regions analysed (**Figures 6A-6C**).

**Figure 6.**
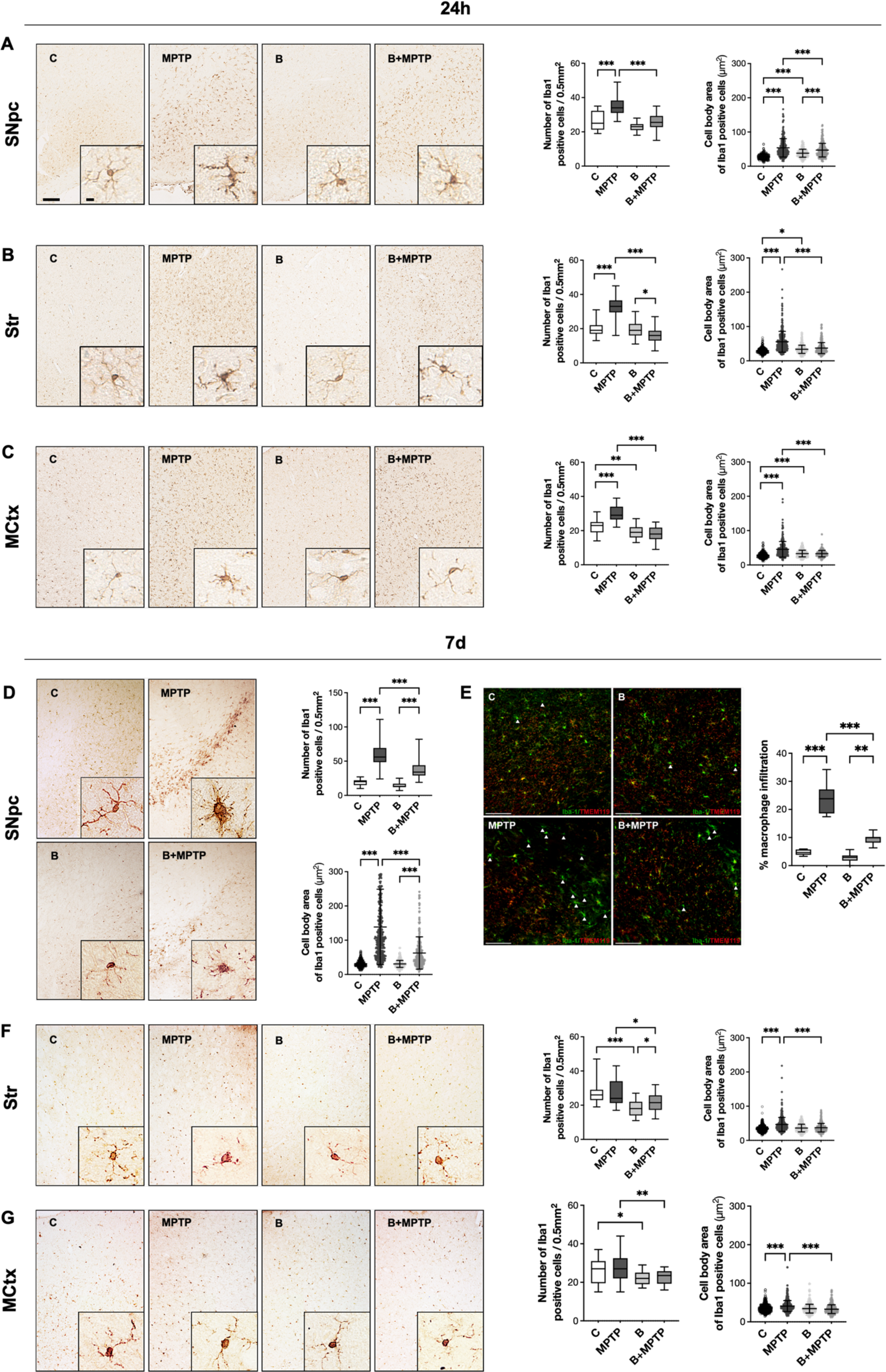
Microglial reactivity based on cell distribution and morphology in the brain 24h and 7d post-MPTP injections, and macrophage infiltration in the SNpc. Number of Iba1-positive (microglial) cells (10x magnification) and cell soma area (40x magnification) in (**A**) Substantia nigra pars compacta (SNpc), (**B**) striatum (Str), and (**C**) motor cortex (MCtx) at the 24h timepoint. Number of Iba1-positive (microglial) cells (10x magnification) and cell soma area (40x magnification) in (**D**) SNpc, (**F**) Str, and (**G**) MCtx at the 24h timepoint. (**E**) Macrophage infiltration in the SNpc at 7d, highlighted by white arrows. Scale bar for 10x magnification 100 µm; for 40x magnification 10 µm. *n*=4 per group; n≥100 cells analysed per animal for body cell area. Statistical significance: *p<0.05, **p<0.01, ***p<0.001. C – Control group; MPTP – MPTP group; B – Berry group; B+MPTP – Berry+MPTP group.

Seven days after the MPTP intoxication, microglial reactivity in the SNpc was more pronounced in comparison with the 24h timepoint (**Figure 6D**). Importantly, at the 7d timepoint, we observed substantial macrophage infiltration in this region following MPTP treatment, which was markedly attenuated in mice receiving the berry-enriched diet (**Figure 6E**). Meanwhile, the number of Iba1-positive cells and their soma size were again significantly reduced in the B+MPTP compared to the MPTP group in the SNpc, striatum, and motor cortex (**Figures 6D, 6F and 6G,** respectively). In the striatum and motor cortex, berry-enriched diet had a strong effect by keeping the number of microglial cells and their soma size at the control levels, parameters which were significantly increased in the MPTP group (**Figures 6D, 6F and 6G**).

### Effect of a berry-enriched diet on astrocyte reactivity is structure and time-dependent

Similar to microglial cells, reactive astrocytes represent propagators of inflammation by releasing proinflammatory cytokines^28^, and their activation has been described to occur after MPTP intoxication^29^. We analysed the GFAP immunostaining intensity to understand whether berry supplementation can affect the activation status of these cells at the two time points studied (**Figures 7A and 7D**). At 24h after MPTP injection, GFAP intensity was not changed in the SN in any group (**Figure 7A**) but was significantly increased in the striatum and motor cortex in both MPTP and B+MPTP groups (**Figures 7B and 7C**), indicating that the berry supplementation did not prevent the activation of astrocytes at this early time point. In contrast, 7d after the MPTP intoxication, we observed a significant increase in GFAP intensity in the SN within MPTP animals and, importantly, a significant reduction in B+MPTP compared to MPTP animals (**Figure 7D**). A similar effect was observed in the striatum, with a clear activation of astrocytes by MPTP that berry supplementation was able to counteract (**Figure 7E**). In the motor cortex, the GFAP intensity was not increased in the MPTP group compared to the control (**Figure 7F**).

**Figure 7.**
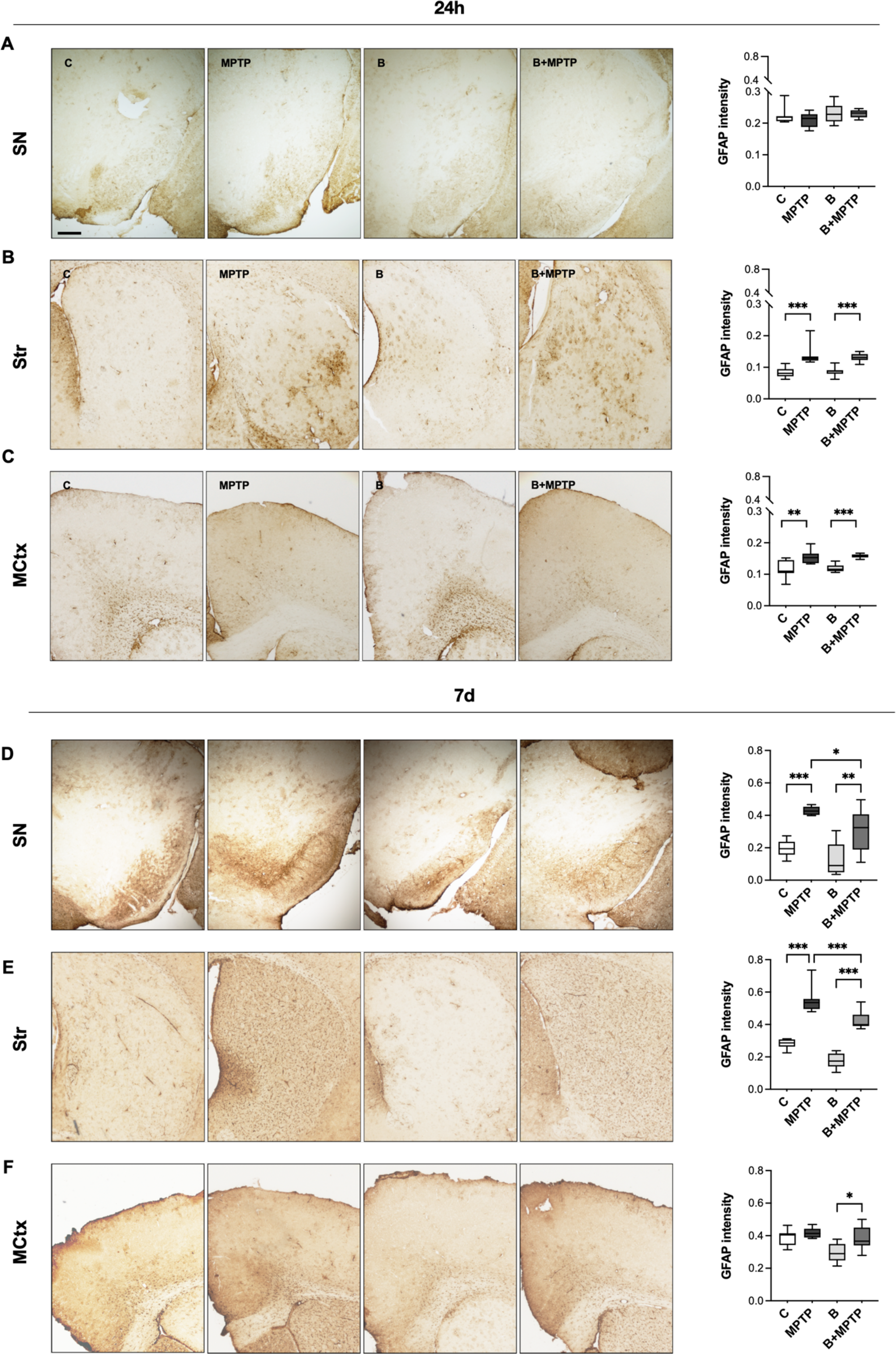
Astrocyte reactivity considering the brain 24h and 7d post-MPTP injections. GFAP staining in (**A**) substantia nigra (SN), (**B**) striatum (Str), and (**C**) motor cortex (MCtx) at 24h post-MPTP injections (5x magnification). GFAP staining in (**D**) SN, (**E**) Str, and (**F**) MCtx at 7d post-MPTP injections (5x magnification). Scale bar: 250 μm; *n*=4 per group. Statistical significance: *p<0.05, **p<0.01, ***p<0.001. C – Control group; MPTP – MPTP group; B – Berry group; B+MPTP – Berry+MPTP group.

### A berry-enriched diet ameliorated inflammatory indicators in serum and brain regions

Following the observed effects of berry supplementation in attenuating signs of neuroinflammation in MPTP-treated mice and given the pivotal role of inflammation in PD pathogenesis, we further investigated inflammation-related transcriptional changes by evaluating cytokine gene expression profiles in the brain that could relate to our findings in microglial cells and astrocytes, both involved in immune and inflammatory responses in the brain. Our results revealed distinct outcomes across the four study groups. Particularly, in the midbrain, *Tnfα* was upregulated at 24 hours post-MPTP injections, an increase that was no longer significant at 7d (**Figure 8A**). This pattern was consistent in the striatum, where *Tnfα* levels were elevated by MPTP at 24h with no differences noted at 7d (**Figure 8B**). The berry-supplemented diet did not prevent the increase in *Tnfα* transcript levels at 24h. *Il1β* mRNA levels in the midbrain exhibited slight upward trends in the MPTP groups at both examined time points, though these changes did not reach statistical significance (**Figure 8C**).

**Figure 8.**
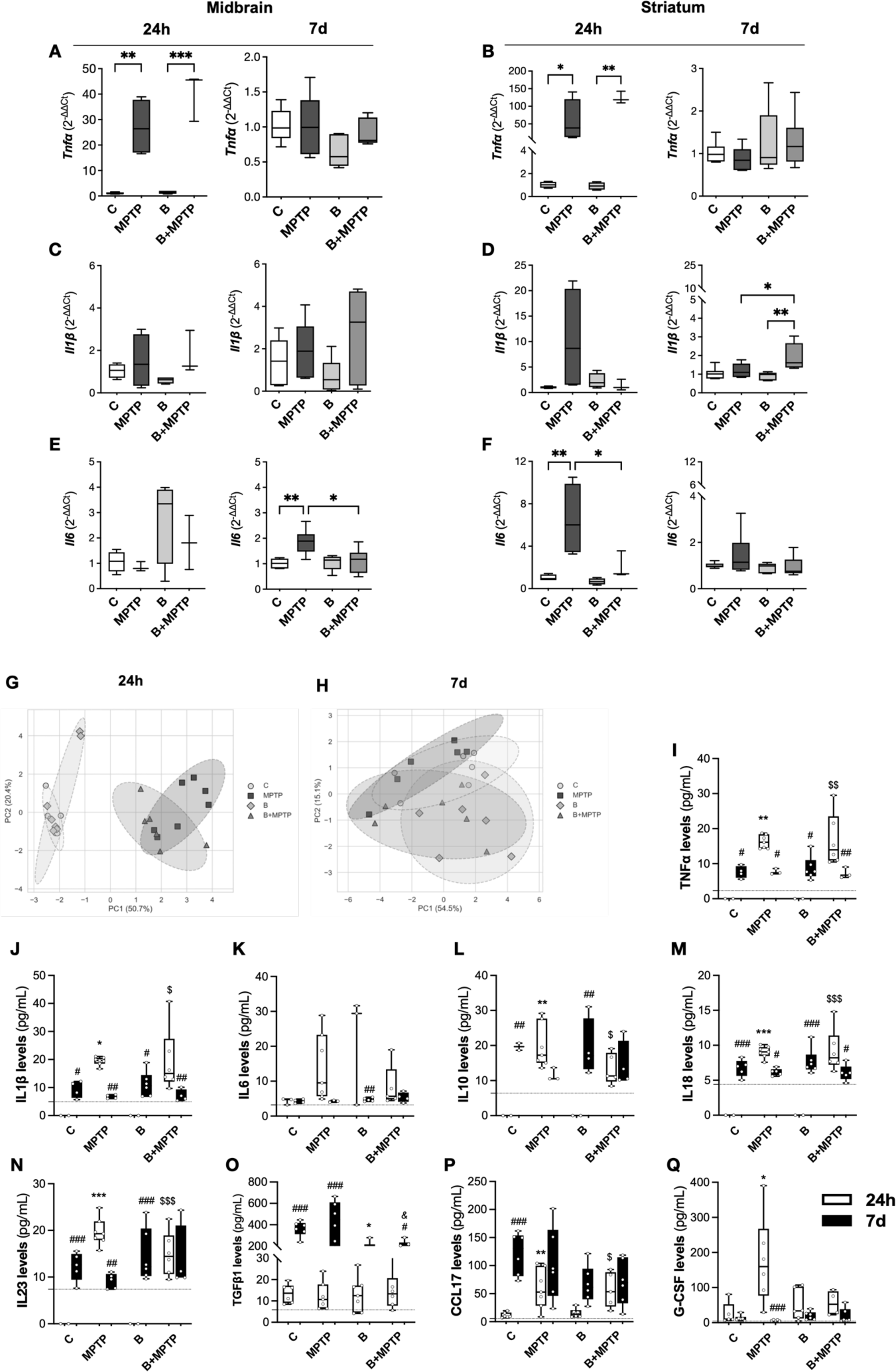
Analysis of cytokine and pro-inflammatory chemokine profiling at 24h and 7d post-injections. mRNA levels of *Tnfα* (**A, B**), *Il1β* (**C, D**), and *Il6* (**E, F**), from the midbrain and striatum at both timepoints. n=3-6 per group. Statistical significance: *p<0.05, **p<0.01, ***p<0.001. Principal Component Analysis (PCA) with the levels of inflammatory-related proteins quantified in serum at (**G**) 24h and (**H**) 7d post-injections. Serum levels of (**I**) TNFα, (**J**) IL1β, (**K**) IL6, (**L**) IL10, (**M**) IL18, (**N**) IL23, (**O**) TGFβ1, (**P**) CCL17 and (**Q**) G-CSF quantified with the bead-based LEGENDplex immunoassay at both timepoints (24h and 7d). Dash lines represent the limit of detection for each molecule. *n*=3-7 per group. Statistical striatum at both timepoints. *n*=3-6 per group. Statistical significance: ^#^p<0.05; ^##^p<0.01, ^###^p<0.01 for same group 24h vs. 7d; *p<0.05, **p<0.01 for comparison with C group; ^$^p<0.05, ^$$^p<0.01 for comparison with B group; ^&^p<0.05: comparison with MPTP group (of the correspondent timepoint). C – Control group; MPTP – MPTP group; B – Berry group; B+MPTP – Berry+MPTP group.

In the striatum, *Il1β* remained unchanged at 24h. However, an unexpected increase in *Il1β* mRNA was observed at 7d in the B+MPTP group (**Figure 8D**). Regarding *Il6*, a significant rise was induced by MPTP in the midbrain at 7d, an upregulation that was mitigated in the B+MPTP group (**Figure 8E**). In the striatum, a similar MPTP-induced increase in *Il6* was noted at 24h, which was prevented in the berry-fed animals (B+MPTP). In contrast, the levels did not differ significantly across the groups at 7d (**Figure 8F**). These findings highlight the complex interplay between MPTP-induced neuroinflammation and the potential modulatory effects of berry supplementation on specific inflammatory markers in the brain.

To complement the brain-specific inflammatory profile, we next explored whether these effects extended to the systemic level by assessing a broader panel of cytokines in the serum, TNFα, IL1β, IL6, IL10, IL18, IL23, TGFβ1, CCL17, G-CSF, CXCL1, and IL12p40. This systemic evaluation aimed to determine whether peripheral inflammatory signals mirrored the central immune response and whether berry supplementation exerted any modulatory effect beyond the brain. A multivariate analysis of serum cytokines by PCA revealed a clear separation between control groups (C and B) and MPTP-challenged groups (MPTP and B+MPTP) at the 24h endpoint (**Figure 8G**). In contrast, the clustering overlapped the different groups after 7d post-injections (**Figure 8H**). In a more detailed analysis, we observed that serum levels of TNFα, IL1β, IL6, IL10, IL18, IL23, CCL17 and G-CSF (**Figures 8I-8N, 8P and 8Q**) were found elevated in MPTP-intoxicated animals after 24h, an increase that berry supplementation could not suppress except for G-CSF (**Figure 8Q**). In particular, TNFα, IL1β, IL10, and IL23 showed no detectable levels in their respective controls at 24h (i.e., below the assay detection limit) (**Figures 8I, 8J, 8L and 8N**, respectively). All remaining cytokines (TGFβ1, CXCL1 and IL12p40) were not significantly altered among the groups after 24h of MPTP treatment (**Figure 8O and Figure S7, Supplementary Information**). After 7d post-injection, TNFα, IL1β, IL6, IL10, IL18, IL23, CCL17 and G-CSF were detected in serum in all four groups, with no differences among them, which may reflect a mild systemic inflammatory response induced by the behavioural testing procedures performed between 48h and 4 days post-injections. (**Figures 8C-8H, 8J and 8K**). IL12p40 and CXCL1 remained unaltered after 7d post-injection (**Supplementary Information, Figure S6**). Of note that the serum levels of TNFα, IL1β, IL6, IL18, IL23 and G-CSF were reduced at 7d compared to 24h, particularly in the MPTP and B+MPTP groups (**Figures 8C-8D, 8G-8H and 8K**). On the contrary, TGFβ1 and CCL17 levels were overall higher at 7d post-injections (**Figures 8I and 8J**).

## Discussion

Neurodegeneration in PD is a process that evolves over time. When the first clinical symptoms occur, the disease is too advanced to reverse the damage already inflicted. Neuroinflammation and oxidative stress have been described as two of the main drivers of neurodegeneration in PD. They appear in the very early phase of pathology, even before symptomatology takes place, and are modifiable^30^, thus presenting a window of opportunity for potential preventive/treatment strategies that can suppress/delay the development of these pathological changes and disrupt the progression of the disease. Therefore, the current worldwide endeavours to alleviate the impact of NDs have focused on filling the lack of effective disease-modifying therapies and developing new preventive strategies. In this regard, diet alterations and consuming (poly)phenol-rich foods, which have been linked to multiple beneficial effects on the brain^31,32^, support the hypothesis that (poly)phenol-rich diets could potentially decrease the risk of developing or delaying the progression PD.

In the current study, we provided evidence that a berry-enriched diet can delay the loss of dopaminergic markers, improve survival of dopaminergic neurons in the SNpc, and ameliorate microglial and astrocytic reactivity in an MPTP-induced mouse model of PD. Importantly, this non-invasive dietary approach successfully mitigated PD-like motor symptoms.

The impairment in motor functions is the most striking symptom of PD that significantly impacts the everyday life of patients. Motor deficits observed in PD are caused by the loss of dopaminergic neurons in the striatum, impairing motor behaviour^33^. To date, no ideal animal model exists that mimics all behavioural, cellular, and molecular aspects of PD. However, the MPTP model is the most widespread and used, as it recapitulates most pathological hallmarks of PD, with motor impairment being one of those^34^. As previously reported in different acute MPTP paradigms, loss of TH-positive neurons in the SNpc, together with a reduction in DA metabolites, like DOPAC and HVA, and impaired motor behaviour were observed^35^. Some preclinical studies employing sub-chronic and chronic MPTP models showed that single (poly)phenols like baicalin and (-)- epigallocatechin-3-gallate, applied either orally or intraperitoneally, exerted neuroprotective effects as they prevented the loss of TH-positive cells in the SNpc and TH levels in the striatum^36–39^. This was accompanied by increased DA, DOPAC and HVA levels in the brain and, consequently, by improved motor behaviour in different regimens of the MPTP intoxication^39–42^. Our study showed that a dietary approach using a berry supplementation significantly protected the loss of TH-positive neurons and undoubtedly improved motor performance in tests requiring fine motor skills and strength, like rotarod, grid, and tight rope test in the acute MPTP model. However, this neuroprotection and motoric improvement was not linked to the increased levels of either DA or its metabolites in berry-supplemented animals. Attending to the neuroprotective effects of berries in alleviating motor dysfunction, we may say that motor performance is not just related to DA levels, and there may be other important mechanisms involved that result in the phenotype observed in the berry-supplemented mice. A very recent study has shown no correlation between the levels of striatal DA and behavioural performance in an acute model of MPTP^43^, pointing to the possibility of (poly)phenols achieving their beneficial effects through some other mechanism(s).

It has been described that higher cortisol levels observed in the serum of PD patients are associated with a higher rate of neurodegeneration^44–46^. A higher rate of neurodegeneration may impact the disease progression since it may trigger or exacerbate motor symptoms^47^. Our biochemical analysis has shown increased levels of corticosterone in the urine of mice upon MPTP intoxication, while berry supplementation completely reverted this increase. Corticosterone release is primarily regulated by the hypothalamic-pituitary-adrenal (HPA) axis, a critical system for maintaining homeostasis and responding to stress. Such a phenomenon is an additional systemic effect of berry supplementation, resulting from a direct effect on the HPA axis or indirectly by mitigating oxidative stress and inflammation, which might explain the observed neuroprotection on motor symptoms.

Throughout decades, (poly)phenols have been mostly explored for their antioxidant properties, improving endogenous enzymatic defences, and reducing lipoperoxidation, among other antioxidant features^48^, potentially crucial in PD management. Although MPTP intoxication did not significantly affect the expression of *Gclc*, *Gclm*, *Nrf2* or *Nqo1* in our study, berry supplementation appeared to influence aspects of the brain thiolome profile. Disrupted redox homeostasis is pivotal in disease progression in neurodegenerative disorders, including PD^11,26,49^. Oxidation of DNA, protein, lipids, and carbohydrates has been associated with ageing and neurodegenerative conditions, which have been linked to the depletion of glutathione and Cys^49,50^. The results obtained from studies on thiols highlight their indispensable function in mitigating oxidative stress as one of the major mechanisms of neuronal death in PD^11,26,49^. The major antioxidants GSH and Cys can neutralise much of the oxidative damage generated. Particularly, Cys is crucial in cellular redox homeostasis, as the availability of Cys is the rate-limiting step for GSH synthesis^49,51^, whose disruption might be the origin of increased oxidative stress, which has been associated with several neurodegenerative disorders^49^. In the present study, we observed a decrease in total Cys, free total Cys, and the CysSSP levels caused by MPTP intoxication 24h post-injection, suggesting an undoubted interference in the thiol-related antioxidant defences. We also observed that the consumption of a berry-rich diet promoted a decrease in total Cys and free total Cys. A similar effect was already described in hypertensive rats, where a berries-enriched diet promoted the metabolic use of Cys, reflected in a decrease in Cys redox pools, as a strategy to better cope with kidney dysfunction in the rats^52^. It has been shown that regulated control of cysteine metabolism is central to optimal neuronal function and that maintaining cysteine balance has therapeutic benefits^49^. It is tempting to speculate that this decrease in Cys levels in the brain just upon berry feeding represents a novel berry-induced Cys balance that is preparing the system, making it more resilient to potential subsequent challenges, and is reflected in unchanged Cys levels upon MPTP intoxication in B+MPTP animals. However, it remains unknown how and why berry supplementation decreases Cys levels in physiological conditions, how it maintains these levels unchanged after the MPTP intoxication and exactly how this affects the neurodegeneration and PD progression. We also observed decreased levels of CysGlySP 24h after the MPTP intoxication, while the berry supplementation managed to keep the CysGlySP levels unchanged even after the MPTP challenge. Very importantly, berry supplementation managed to make a new homeostasis of cysteine-related thiolome, making it stable even upon very harsh challenges like MPTP intoxication. Nevertheless, further research is needed to uncover how berry supplementation interferes with cysteine-related thiolome and its effects against neural insults observed in neurodegenerative diseases.

Neuroinflammation, reflected in elevated glial reactivity, represents one of the main processes underlying PD pathology^5,28^. Anti-inflammatory treatment in PD patients is aimed to tackle this process, and in fact, recent studies suggest that common nonsteroidal anti-inflammatory drugs (NSAIDs) like ibuprofen may have a protective effect in individuals with PD in delaying the average delay of 8.2 years in the onset of Parkinson’s symptoms compared to those who had not^53^. However, long-term use of NSAIDs is not an option in clinical practice for these patients since it can lead to serious side effects, including gastrointestinal bleeding, kidney and liver damage, and increased cardiovascular risk. Nevertheless, attenuation of neuroinflammation is a strategy that needs to be exploited for PD management. In fact, several preclinical studies have shown that just suppressing neuroinflammation is enough to obtain neuroprotection and improved motor behaviour in PD models^30,54,55^. Previous studies exploring the effects of (poly)phenols in *in vitro* and *in vivo* PD models undoubtedly showed that these compounds exert neuroprotective and anti-inflammatory properties^16,42,56–58^. However, most of these studies explored a single pure (poly)phenol, very often as a pre-treatment, at high concentrations and frequently through the intraperitoneal route, pointing to a pharmacological approach. Nonetheless, administering medication prior to diagnosis is not a standard clinical practice, and the use of pure (poly)phenols as a drug for prolonged periods will pose safety implications due to high concentrations of compounds. Including a preventive approach through diet can be very attractive for people at risk of developing PD, and on the other hand, it can be a complementary approach to medication after PD diagnosis to delay the progression of the disease. In fact, neuroinflammation is an early symptom that appears in PD pathology years before the occurrence of the first symptoms^17^. Our study explores this different paradigm based on the effect of a healthy (poly)phenol-enriched diet as a preventive approach for individuals at risk of developing PD and a strategy to delay progression for early stages of a PD patient by attenuating systemic and neuroinflammation. We observed a significant increase in microglial cell number and cell body size after the MPTP injections, which aligns with previous studies that show a widespread switch in microglial phenotype upon MPTP intoxication^13,29,59^. Noteworthy, our dietary approach consisting of berry supplementation was very effective in mitigating the neuroinflammation induced by MPTP, as evidenced by the decrease in both the number and morphology of microglial cells across all three targeted brain regions (SNpc, striatum and motor cortex), both 24h and 7d timepoints. On the other hand, the beneficial impact of berry supplementation on astrogliosis was also observed at 7d in SN and striatum, while no effect was observed at 24h post-MPTP exposure, suggesting that berries primarily modulate microglial rather than astrocytic inflammation. Another very important finding of our study is that microglial soma size was increased in the B group in all three brain regions at 24h timepoint, just upon berry feeding, stressing the impact of berry supplementation on microglial cells without any challenge, suggesting microglia activation. This phenomenon suggests a pre-conditioning effect of the cells that might help microglia cells to deal with subsequent stress conditions. The MPTP-induced microglia morphology alterations were significantly alleviated in berry-supplemented animals in all three brain regions after the MPTP intoxication, favouring our hypothesis. Concomitant with glial cells activation, the release of inflammatory cytokines aggravates the degeneration of dopaminergic neurons in the SN^60^. In our study, we observed increased TNFα expression levels in both the midbrain and striatum at the early 24h timepoint after MPTP injections, which were not prevented by berry supplementation. MPTP increased *Il6* mRNA levels in the brain, an effect that the berry-supplemented diet was able to prevent at both 24h and 7d after MPTP administration. The upregulation of *Il6* has been shown to exacerbate dopaminergic degeneration in 6-hydroxydopamine-induced PD rats^61^, while a four-year prospective study also demonstrated that IL6, but not TNFα, contributes to mortality in PD patients^62^. Therefore, IL6 is crucial to neuroinflammation and has emerged as a pivotal player in PD. Although some previous studies showed that treatment with (poly)phenols can reduce brain levels of IL6 in mice after the acute MPTP intoxication, these studies were done using a single pure (poly)phenol, resveratrol^63^, or an açai extract^21^ at very high concentrations, referring more to pharmaceutical/nutraceutical than dietary approach. Also, most studies exploring the anti-inflammatory potential of (poly)phenols in PD models have focused on analysing TNFα and IL1β and neglected IL6^15^. To the best of our knowledge, our study is the first nutritional study using a (poly)phenol-rich food in the MPTP model of PD, showing that this approach might exert its effects on neuroinflammation through the inhibition of IL6 expression rather than TNFα and IL1β. A very elegant study done by Cardenas and Bolin^64^, using IL6 knockout mice (^−^/^−^) and acute MPTP intoxication, showed that while astrogliosis was similar in the SNpc of IL6 (^+^/^+^) and IL6 (^−^/^−^) mice upon MPTP, microgliosis was severely diminished in IL6 (^−^/^−^) mice. They showed that in the absence of IL6, an acute reactive microgliosis was transient, with a complete absence of reactive microglia at day 7 post-MPTP intoxication. In contrast, extensive reactive microgliosis was observed in the SNpc of MPTP-intoxicated IL6 (^+^/^+^) mice. In our study, similar levels of astrogliosis were observed in both MPTP and B+MPTP animals at 24h in the striatum and motor cortex, as well as at 7d in the motor cortex and this is concomitant with a significant reduction in microglia activation by berry supplementation. This reduction is reflected in both microglial cell numbers and morphological changes in the same tissues. Additionally, our findings point to the brain (poly)phenol-induced IL6 expression decrease as a possible mechanism underlying the anti-neuroinflammatory potential of our dietary approach.

One striking finding is that IL6 was increased at the 24h timepoint, although not significantly, in the midbrain of the B group, which was related to a significantly increased microglia soma size in the SNpc, just upon berry supplementation. Recently, IL6 emerged as an important regulator of neuronal and glial function in health and disease, being described as a pleiotropic cytokine, which might exert neuroprotective or neurotoxic effects depending on the context^65^. Therefore, it is tempting to suggest that the observed slight increase in IL6 expression in the midbrain, just upon berry feeding, might serve as a protective mechanism triggered by berries to prepare the system for the insult. Since the berry supplementation sustained the expression of IL6 upon MPTP insult at the control level in the midbrain, it suggests that the mild increase observed just with berries may give an advantage to these animals compared to the control in response to MPTP. However, further studies are needed to fully understand how and why berry feeding induces this effect and its physiological meaning.

Besides inflammatory response in the brain, peripheral inflammation is increasingly believed to play an important role in PD pathogenesis^66^. Evidence suggests that immune dysregulation in both the periphery and CNS contributes to the upregulation of inflammatory mediators^67^. In line with the predictable dynamic changes triggered by acute MPTP exposure^29,68^, our study showed a distinct profile with increased levels of inflammatory cytokines in the serum of MPTP-intoxicated mice at 24h post-injection, a timepoint often associated with the peak of inflammatory response^29^. By 7 days post-MPTP, systemic cytokine levels had largely returned to baseline, suggesting inflammatory resolution. However, at this same 7d timepoint, we observed pronounced macrophage infiltration in the SNpc, supporting the notion that peripheral immune cells can access the brain parenchyma in response to neurodegeneration^69^. This is consistent with data showing that BBB disruption facilitates peripheral immune cell entry and that MPTP exposure leads to increased leakage^70^. Notably, this infiltration was markedly attenuated by the berry-enriched diet, supporting its potential to modulate not only central glial responses but also immune cell trafficking from the periphery. Recent data have gathered insights that LMWPMs can preserve BBB function and reduce immune cell trafficking in neurodegenerative cellular models^71–73^. The increased presence of TNFα, IL1β, IL10, IL18, IL23, CCL17, and G-CSF observed in the serum of MPTP animals after 24h agrees with the increased pro-inflammatory cytokines and chemokines found elevated in blood and CSF from PD patients^74–76^. Increased levels of TNFα, IL1β, and IL6 in the serum of PD patients have been shown to correlate with disease severity and motor disability^76^. Berries supplementation before MPTP insult could not sustain the levels of these cytokines. Similarly to what we have observed in the brain, TNFα and IL1β were significantly elevated in 24h and unchanged in 7d after the MPTP intoxication. IL6 followed the same pattern as in the brain by being slightly induced at 24h by MPTP and upon berry feeding itself. Although this difference did not reach statistical significance, the obvious trend observed with serum IL6 and the alterations of the same cytokine in the brain sheds light on IL6 as a possible mediator of berry (poly) phenols’ action in alleviating neuroinflammation and neurodegeneration.

Importantly, although (poly)phenols as potential effectors for NDs have been in the spotlight in past years, and their positive effects on the brain have been suggested, their effects after a nutritional intervention in a disease model and what are the specific metabolites reaching the brain and their concentrations in this scenario remain to be defined. Dietary (poly)phenols exhibit limited bioavailability, attributed to interactions with the food matrix, metabolic processes facilitated by the liver and/or the intestine (involving phase I and II metabolism), and the gut microbiota metabolism^18^. Therefore, the compounds that ultimately reach circulation and organs are predominantly LMWPMs rather than the original phenolic parent compounds^19,20^. We and others showed that LMWPMs appear in circulation, being able to cross the BBB and reach the brain, supporting the hypotheses on direct effects of (poly)phenols on brain function^77^, making them very appealing as potential bioactive compounds in both preventive and therapeutic settings^78–81^. We showed that following a 6-week feeding of a berry-enriched diet, a wide range of (poly)phenol metabolites increased in circulation in C57BL/6J mice. The substantial number of circulating metabolites detected in mice, despite the absence of their parent compounds in the standard diet, may reflect the increased sensitivity of plasma analyses, which enables the detection of a broader spectrum of metabolites, including those arising from extensive biotransformation processes such as hydrolysis. Moreover, we found some unique metabolites detected exclusively in the plasma of berry-fed animals, such as 5-(3’,4’-dihydroxyphenyl)-*γ*-valerolactone sulfate, 5-(phenyl)-*γ*-valerolactone sulfate, and Methyl-*O*-epicatechin-sulfate, clear indicators of flavan-3-ols from the (poly)phenol-enriched diet consumption.

Interestingly, considering the mouse brain, our analysis revealed significant differences in the metabolomic profile between the control and berry groups, establishing distinct baseline conditions before MPTP-induced toxicity.

We observed a selective increase in certain metabolites following consumption of a berry-enriched diet, like pyrogallol, phloroglucinol, and caffeic acid, while some were decreased, like 3-(4′-hydroxyphenyl)propanoic acid, dihydrocaffeic acid, dihydroxybenzaldehyde, and phloroglucinaldehyde, compared to the controls, a differential brain metabolite content that potentially could be related to the observed preventive effects. It is of notice that these are persistent metabolites, which means the metabolites that are still detectable in the brain at the time of sacrifice. We may consider that metabolites present in the brain are transient, like in circulation, since they are further metabolised to be eliminated. Furthermore, it should be considered that MPTP may influence the biotransformation of these metabolites since MPTP is described to alter the expression and activity of CYP enzymes, largely involved in phase I metabolism, potentially affecting the metabolism of their substrates^82^, including (poly)phenols. The significant increase observed in the brain for phloroglucinol and pyrogallol in the animals fed with berries was not registered following MPTP administration.

Concerning pyrogallol, it is important to highlight that it was the most abundant metabolite in the brain of the animals. The concentration highlights the brain permeability and penetrance, as already described for pyrogallol and other LMWPMs^73,81^. Remarkably, pyrogallol sulfate, has been described as very active in attenuating various hallmarks of NDs, in particular, by improving neuronal cell responses to oxidation, excitotoxicity, and even a strong attenuator of microglia activation at physiological levels^83^. Additionally, pyrogallol sulfate and catechol sulfate protected dopaminergic neurons in a 3D cellular model from MPP^+^ insult. Interestingly, in this PD cellular model, the metabolites pre-conditioned dopaminergic neurons by triggering molecular mechanisms that help neurons cope with a stronger and subsequent toxic insult ^84^. Previously, we have described that in human brain microvascular endothelial cells (HBMEC), pyrogallol sulfate is converted to pyrogallol, suggesting the presence of arylsulfatase in those cells^83^. Thus, it is possible that pyrogallol-sulfate in the brain has been converted to pyrogallol. Importantly, pyrogallol, phloroglucinol and caffeic acid, which were significantly increased in the brains of the berry group, were already reported to reach the CSF of individuals taking a purified anthocyanin supplement for 24 weeks^85^. This points out the potential translation of our data for Parkinson’s patients.

Overall, herein we showed that a dietary approach consisting of a (poly)phenol-rich diet can substantially modulate neuroinflammation, oxidative stress, and neurodegeneration, improving motor symptoms of PD. Moreover, we discovered that berry supplementation induced changes in some neuroinflammatory and oxidative stress markers, leading to a better response after the MPTP intoxication, pointing to a putative pre-conditioning effect of berries. Most importantly, we revealed the fingerprint of (poly)phenols and derived metabolites occurring in the circulation and brains of berry-supplemented animals, which might be some of the real effectors behind all these positive effects observed in this study. To the best of our knowledge, this is the first study applying a purely dietary approach consisting of a (poly)phenol-rich food in the MPTP PD-like pathology, showing that a simple, non-invasive, and sustainable change in everyday life can substantially affect PD pathology at various levels. Given the current lack of disease-modifying therapies for PD, any approach that may delay the onset of PD, and therefore prevent years of disability will be instrumental. This work is one of the first rock-solid information on the nutritional value of (poly)phenols for PD that may help in the future translation of these conclusions to humans. Furthermore, the huge potential of (poly)phenols highlights their dual value as both a dietary intervention and a foundation for drug development, making them a significant class of compounds to consider in pharmacological research.

## Materials and Methods

### Animals and experimental design

For the experiment, 4-week-old male C57BL/6J mice were purchased from Charles River (Charles River, France). The animals were housed in pairs before entering the study, submitted to a one-week quarantine period, followed by an additional week for acclimatisation. At this stage, animals were randomly assigned into four groups and maintained in a humidity-controlled room under a 12h light/dark cycle, 55% relative humidity at 20–22°C, and allowed to consume food and water *ad libitum*. At 6 weeks of age, animals were randomly divided into four groups. Two groups were given a standard diet (Teklad 2018, Envigo, US), whereas the remaining two groups were fed with an 8% berry-enriched-standard diet, composed of a lyophilised mixture of blueberries (*Vaccinium* spp. variety Star), blackberries (*Rubus* spp. L. variety Loch Ness), and raspberries (*Rubus idaeus* L. variety Kweli) produced in Odemira region, Portugal. Food intake was monitored daily, and animals’ weight was controlled weekly throughout the trial. PD-like phenotype was achieved by administering an acute dose of MPTP hydrochloride (Sigma) to 12-week-old male mice. Mice from each diet regimen were administered intraperitoneal injection either with 4x15 mg MPTP/kg of body weight dissolved in 0.9% saline (**MPTP** group – MPTP-treated animals fed with standard diet; **B+MPTP** group – MPTP treated mice fed with berry-enriched diet) or only saline (**C** – standard diet-fed control group; **B** – berry-fed group) at 2h intervals. Mice of all four groups were sacrificed either at early (24h) and late (7d) timepoints after MPTP (**Figure 1A**). All procedures were carried out following the guidelines and regulations under Direção Geral de Alimentação e Veterinária (DGAV) approved license 0421/000/000/2021 by researchers accredited by the Federation of European Laboratory Animal Science Associations (FELASA)/DGAV.

### Biological samples

Animals were weighed and overdosed with the volatile anaesthetic isoflurane (IsoFlo, 100% Isoflurane, ECUPHAR). Mice were perfused with ice-cold saline. A total of *n*=8-10 or *n*=14-16 animals per each of the four experimental groups were sacrificed after 24h or 7d, respectively. Blood and urine samples were collected from the right atrium and bladder before perfusion. Perfused brains were isolated, and both hemispheres were divided. The right side was immediately placed in 4% paraformaldehyde (PFA) for immunohistochemical analysis, and the left side was assigned for cortex, midbrain, and striatum dissection, frozen in liquid nitrogen and stored at -80°C.

### UPLC-ESI-QTOF-MS analysis of brain tissue and plasma samples

Blood and perfused brain samples from the 7d timepoint were used for metabolomic analysis. The plasma was obtained after centrifugating the blood (3,000 × g for 10 min at 4°C) and kept at -80°C. Plasma samples (100 µL) were extracted with a mixture of 600 µL acetonitrile:formic acid (98:2 v/v) following the protocol as previously described^86^. From each brain sample, 200 mg was weighed and rinsed with cold phosphate-buffered saline (PBS). Stainless steel beads were added to each sample, and a mixture of 1.25 mL of methanol:hydrochloric acid (99.9:0.1 v/v) was added for the metabolite extraction. Brains were homogenised using a Bullet Blender Homogeniser (Next Advance, Averill Park, NY, USA) for 5 min. After the breakdown of the tissues, samples were sonicated in an ultrasonic bath for 10 min and centrifuged at 10,000 × g for 10 min at 4°C. Each supernatant was evaporated in a speed vacuum, re-suspended in 100 μL of methanol, and filtered through a 0.22 μm filter.

Analyses of (poly)phenol metabolites present in the brain tissue and plasma samples were performed on an Agilent 1290 Infinity UPLC system coupled to a 6550 Accurate-Mass QTOF-MS (Agilent Technologies, Waldbronn, Germany) using an electrospray interface (Jet Stream Technology)^86^. A target screening strategy^86^ was applied to qualitatively analyse 132 metabolites potentially present after berry consumption (**Supplementary Information, Table S4**). These compounds included conjugated metabolites (e.g., glucuronides and sulfates). The exact mass of each proposed compound was extracted using an extraction window of 0.01 m/z.

### Corticosterone

Corticosterone levels were quantified from the mouse urine samples using the Corticosterone Competitive Enzyme-Linked Immunosorbent Assay (ELISA) Kit (EIACORT, Invitrogen) according to the manufacturer’s instructions.

### Behavioural testing

The motor function of MPTP-induced mice was evaluated using six behavioural tests performed on all animals, from day 2 to day 5 post-MPTP injections (Figure 1A). Open field test to assess mice’s anxiety, motor activity, and exploration^87^ was performed in a 40×40×40 cm acrylic box. Each mouse was placed in the centre and allowed to explore for 20 min. Locomotion videos were analysed with Bonsai Software^88^, recording latency to leave the centre, entries into the inner field, time spent in different areas (inner field, near the wall), ambulatory distance, ambulatory and resting time, and travel speed. The rotarod test evaluated motor coordination, balance, and endurance^89^ using an automatic rotarod Series 8 apparatus from IITC Inc. Life Scienc (CA, USA). Mice were introduced to the test at 7 rpm for 15 min to learn how to perform the test. The apparatus then accelerated from 4 to 40 rpm over 300 s. Latency to fall was recorded over three trials per mouse, with 15-min intervals. Results were averaged and normalised to the baseline. Kondziela’s inverted screen test to assess muscular strength, motor performance, and coordination^90,91^ was done by placing mice on a metal grid 50 cm above a padded surface, which was then inverted. The test lasted 3 min and was repeated 3 times with 15 min intervals. The time taken to fall was recorded. Mice falling before 3 min scored 0, while those remaining for 3 min scored 1. The average of the three trials was used. The global muscle function, coordination, and balance were assessed by the tight rope test^92^. It uses a 1 m long, 3 mm thick steel wire covered with soft plastic, stretched 60 cm above the ground between two platforms. Mice are trained to grab the rope and return to the platform. During trials, mice are placed in the centre and given 60 sec to reach the platform. Scores are based on performance: 3 for reaching the platform, 2 for staying on the rope, and 1 for falling. Three trials are conducted, and the average score and time are calculated. Finally, to assess locomotor activity, motor coordination, and balance, the pole test was performed^93,94^. It uses a 60 cm wooden pole with a 1 cm diameter, wrapped in paper tape, placed in a cage with bedding. Mice are trained to descend the pole head-downward. Four trials per mouse are performed, with 15-min intervals. Trials are video-recorded, and times to turn, climb down, and total descent are measured. Data from the four trials are averaged and presented with standard deviation.

### Western Blotting

Frozen samples used in western blotting assays were homogenised in cell lysis buffer from Cell Signalling supplemented with 0.1 mM phenylmethylsulfonyl fluoride (PMSF), 2 mM dithiothreitol (DTT), and protease and phosphatase inhibitor cocktails (PhosSTOPTM and cOmpleteTM, EDTA-free Protease Inhibitor, respectively). The resulting homogenates were centrifuged at 16,100 × g for 10 min at 4°C, and the supernatants were stored at -80°C until use.

The protein concentration of each sample was quantified using the bicinchoninic acid (BCA) protein assay kit, following the manufacturer’s instructions. The samples followed a denaturation protocol by adding 4x concentrated sample buffer (0.24 M Tris-HCl, 8% sodium dodecyl sulfate (SDS), 40% glycerol, 0.6 M DTT, 0.1% bromophenol blue; pH 6.8) and heating for 5 min at 95°C. Samples were then separated in sodium dodecyl sulfate-polyacrylamide gel electrophoresis (SDS-PAGE) and electro-transferred onto a PVDF membrane. Nonspecific binding was blocked by incubating the membranes with 5% bovine serum albumin (BSA). Membranes were incubated with primary antibodies against TH (1:500, #SC-14007, Santa Cruz Biotechnology), Bcl-2 (#AB0378-200, Sigma), Bax (#AB0164-200, Sigma), and GAPDH (#MA5-15738, Invitrogen). After rinsed in Tris-buffered saline buffer (137 mM NaCl, 20 mM Tris–HCl; pH 7.6) containing 0.1% Tween-20 (TBS-T), the membranes were incubated at RT in Horseradish Peroxidase (HRP)-or Alkaline Phosphatase (AP)-conjugated secondary antibodies diluted in 5% BSA, for 1h. The detection was performed using the Amersham ECL Prime Detection Reagent, and visualisation was carried out with the Odyssey® Fc Dual-Mode Imaging System. Band intensity was digitally quantified using Image Studio™ software.

### Quantification of cysteine-related thiolome

The cysteine-related thiolome was obtained through the analysis of the amino acid Cys, the dipeptide cysteinylglycine (CysGly) and the tripeptide GSH. These molecules were analysed in their non-protein-bound forms, comprising the reduced (RSH) and the respective disulfides (RSSR) fractions, and in their *S*-thiolated protein (RSSP) forms (CysSSP, GSSP and CysGlySSP, respectively). The quantification was obtained by HPLC (Shimadzu Scientific Instruments Inc., Columbia, MD, USA) with an RF 10AXL fluorescence detector as previously described^84,95^. These molecules were separated in a reversed-Phase C18 LiChroCART 250-4 column (LiChrospher 100 RP-18, 5 μm, VWR, Radnor, PA, USA) in a column oven at 29°C on isocratic elution mode for 20 min, at a flow rate of 0.8 mL/min. The mobile phase was constituted by a 100 mM sodium acetate buffer (pH 4.5) with methanol (99:1 (v/v)). The detection was performed using excitation and emission wavelengths of 385 and 515 nm, respectively. Also, the protein-bound fraction of each moiety was calculated by subtracting the total free fraction from the total fraction.

### Neurotransmitters UPLC-ESI-QqQ analysis

Frozen striatum samples used in UPLC-ESI-triple Quadrupole (QqQ) analysis were weighted and homogenized with 400 µL of a cold mixture of acetonitrile:formic acid (98:2, v/v) in a Bullet Blender Homogeniser for 5 min. Afterwards, samples were centrifuged at 14,000 × g for 5 min and the supernatants were evaporated in a speed vacuum. Finally, evaporated samples were re-suspended in 100 μL of the diluting solution containing water:acetonitrile:formic acid:ascorbic acid (96.9:3:0.2:0.02) and filtered through 0.22 μm RC.

MS spectra were acquired in negative or positive ionisation mode (depending on the metabolite). Water:formic acid (99.9:0.1, v/v) and acetonitrile:formic acid (99.9:0.1, v/v) were used as mobile phases A and B, respectively, with a flow rate of 0.4 mL/min. The following gradient was used: 0 min 0% B; 0–6 min 20% B, 6–6.1 min, 20–90% B and 6.1–8 min 90% B. Finally, the B content was decreased to the initial conditions (0%) in 10 sec, and the column was re-equilibrated for 2 min. All compounds were monitored in the multiple reaction monitoring mode. Two transitions with the highest intensity for each compound: a quantifier and at least one qualifier, were optimised, establishing a fragmentor voltage and collision energy for each transition. The optimisation was developed by infusing 50 μM of each compound into the mass spectrometer with a mixture 50:50 of phases A and B at a flow rate of 0.2 mL/min. In parallel, DA, HVA, DOPAC, 5-HT, NE, and ACh calibration curves were prepared freshly by diluting the stock (diluted in water) in the diluting solution. The volume of injection in the equipment was 15 µL.

### Real-time PCR

The total RNA was extracted from 30 mg of frozen tissue, previously disrupted and ground, using the RNeasy Mini Kit (Qiagen) according to the manufacturer’s instructions. 200 μg of total RNA was used for a reverse transcription reaction (total volume 20 µL) using SuperScript^™^ II Reverse Transcriptase (Invitrogen, Carlsbad, CA) and random primers (Roche). Real-time PCR was performed using a TaqMan Fast Advanced Master Mix (Applied Biosystems). The cycle parameters were as follows: Uracil-DNA glycosylase incubation at 50°C for 2 min, polymerase activation at 95°C for 10 min, denaturation at 95°C for 15 sec, and then annealing and extension at 60°C for 1 min. Gene expression assay for NAD(P)H quinone dehydrogenase 1 (*NQO1*, Mm01253561_m1), Glutamate-cysteine ligase catalytic subunit (*Gclc*, Mm00802658_m1), Glutamate-cysteine ligase modifier subunit (*Gclm*, Mm01324400_m1), Nuclear factor erythroid 2– related factor 2 (*Nfe2I2* (Nrf2), Mm00477784_m1), Tumor necrosis factor-α (*Tnf*α, Mm00443260_g1), Interleukin-6 (*Il6*, Mm00446190_m1), Interleukin-1β (*Il1β*, Mm00434228_m1), *Gapdh* (Mm99999915_g1) and Actin (*Actb*, Mm00607939_s1) were from Thermo Fisher. Two endogenous control genes, *GAPDH* and *Actb*, were used for normalisation. The comparative cycle threshold (CT) method was used to calculate the relative amount of transcripts in all groups, and genes were normalised to the endogenous controls. The formula is as follows: ΔΔCT= ΔCT_sample_−ΔCT_normal_, where ΔCT is the difference in CT between the targeted gene and housekeeping controls by minimising the average CT of the controls. The fold-change was calculated as: 2^-ΔΔCT^.

### Immunostaining analysis

For immunostaining analysis, mouse brain halves immersed in 4% PFA for 24h at 4°C were then cryoprotected with incubations in three rounds of 30% sucrose solutions in 0.01 M PBS (for 48h at 4°C each). The brains were frozen in isopentane, cooled on dry ice and stored at -80°C. Every third coronal section (20 µm thick) through the motor cortex and striatum level and every coronal section (20 µm thick) through the midbrain level was mounted on slides, dried overnight and stored at -20°C. For all further analyses, a total of three sections throughout the midbrain (with 400 µm intervals between each section) were used from each brain. A total of 4 brains were used for each experimental group. A total of six sections per animal (separated by 400 µm interval for the motor cortex and the striatum and 200 µm interval for the SNpc) were analysed. For glial cell labelling, Iba1 immunostaining was used to detect microglia and macrophages, while GFAP immunostaining was performed to identify astrocytes. Additionally, we employed a complementary experiment to distinguish resident microglial cells from infiltrating macrophages. Sections were incubated with anti-Iba1 (1:500, #019-19741, Wako), anti-GFAP (1:2000, #Z0334, DAKO) or Anti-TMEM119 (1:2000, #ab209064, Abcam) primary antibodies, followed by the EnVision® System-HRP Labelled Polymer Anti-Rabbit secondary antibody (#K4003, DAKO, Denmark). Imaging was performed using a Zeiss Axio Imager microscope, with 10x and 40x objectives for microglia and 5x for astrocytes. Quantification was carried out using NIH ImageJ 1.50i (National Institutes of Health, Bethesda, MD, USA). To quantify TH-positive neurons in the SNpc, TH/MAP2 double immunostaining was conducted using primary antibodies against TH (1:250, #2928, Sigma Aldrich) and MAP2 (1:1000, #ab5392, Abcam). Secondary antibodies included Alexa Fluor Goat anti-mouse 555 (#A-21422) and Alexa Fluor Goat anti-chicken (#A-11039, Invitrogen). Images were obtained using a Zeiss LSM 710 confocal point-scanning microscope (Zeiss, Germany) with a 20x objective, and analysis was performed using NIH ImageJ 1.50i. Further details in **Supplementary Information** – **Methods Section**.

### Cytokine quantification by flow cytometry analysis

A bead-based multiplex assay was employed to quantify cytokines (LEGENDplex^TM^ Mouse Macrophage/Microglia Panel (13-plex) with V-bottom Plate (BioLegend Inc., USA) in serum samples obtained from the mice, according to the manufacturer’s instructions. Briefly, blood samples were allowed to clot for at least 30 min following centrifugation for 20 min at 1,000 × g. The serum was recovered in the supernatant and stored at -20°C until use. For the assay, 25 μL of 2-fold diluted serum samples, diluted standards, and blanks were added to the plate; 25 μL of premixed beads and detection antibodies were added to all the samples. The plate was incubated for 2h at RT with shaking in the dark. 25 μL of streptavidin–phycoerythrin conjugate was added for 30 min at RT. Then, the samples were washed and suspended in 200 μL of wash buffer. The data were acquired with BD FACS Canto™ II Flow Cytometry System. Bead excitation was achieved using 488 and 640 nm lasers, and the emission was detected using 530/30 and 665/20 nm bandpass filters, respectively. The data were processed with the BioLegend LEGENDplex Data Analysis Software. The concentration was expressed as pg/mL.

### Statistical analysis

Results are presented using box-and-whisker plots. Boxplots represent the median as the middle line, with the lower and upper edges of the boxes representing the 25 and 75% quartiles, respectively; the whiskers represent the range of the full data set, excluding outliers. Statistical analysis was performed using the GraphPad Prism® 9.5.1 software (GraphPad Software, USA). One-way ANOVA was used to compare differences among conditions and groups, followed by Dunnett’s multiple comparison test. The Kruskal–Wallis test was applied for non-parametric data with an abnormal distribution. Statistically significant differences were considered when *p*<0.05.

Multivariate statistical analysis was carried out on the peak intensities for both plasma and brain metabolome using MetaboAnalyst (https://www.metaboanalyst.ca). Data were subsequently analysed by Principal Component Analysis (PCA) to detect intrinsic clusters and outliers within the data. Features (metabolites) were filtered based on their relative standard deviation (RSD). Any feature with an RSD higher than 25% was removed from the analysis to reduce variability and focus on more consistent data.

## Acknowledgements and Funding

This work was supported by the European Research Council (ERC) - Grant No. 804229 and the European Union under Horizon Europe - Grant No. 101060346). Research Units: UID/04462, iNOVA4Health – Programme in Translational Medicine; iMed (UIDB/04138/2020 and UIPD/04138/2020), financially supported by Fundação para a Ciência e Tecnologia / Ministério da Educação, Ciência e Inovação. Programa Regional de Fomento de la Investigación Científica y Técnica (Plan de Actuación 2022) de la Fundación Séneca-Agencia de Ciencia y Tecnología de la Región de Murcia, Spain (22030/PI/22), and PID2022-136419OB-I00, funded by MCIN/AEI/10.13039/501100011033 and “ERDF A way of making Europe” by the European Union. The authors would like to acknowledge FCT for the financial support of R.C. (PD/BD/135492/2018), C.P. (2023.00453.BD), C.P. (2022.11465.BD) and D.C. (2020.04630.BD), Fátima Martins and Inês Silva for their technical support with the endpoint experiments, and to Pedro Oliveira for providing berry fruits.

## Author contributions

R.C., N.L.V. and C.N.S. contributed to experimental design and interpretation. R.C., A.F.R., H.T.L., C.P. and N.L.V. performed experiments. M.A.A.G., C.O.S., A.M. performed HPLC-related experiments. S.A.P., A.G.S., J.C.E., A.N.C, D.C, T.F.P. and C.N.S. provided materials and valuable knowledge. R.C., N.L.V. and C.N.S. wrote the manuscript with feedback and input from all authors. All authors read and approved the final manuscript.

## Additional information

Supplementary information: The online version contains supplementary material available at

## Data availability

The data analysed during this study are available from the corresponding author upon request.

