## Supplemental information for "From berries to brain: Assessing the impact of (poly)phenols in the MPTP mouse model of Parkinson’s disease"

**1** iNOVA4Health, NOVA Medical School, Faculdade de Ciências Médicas, Universidade NOVA de Lisboa, Lisboa, Portugal.
**2** Instituto de Tecnologia Química e Biológica António Xavier, Universidade NOVA de Lisboa, Oeiras, Portugal.
**3** Laboratory of Food & Health, Research Group on Quality, Safety, and Bioactivity of Plant Foods, Campus de Espinardo, CEBAS-CSIC, Murcia, Spain.
**4** Research Institute for Medicines (iMed.ULisboa), Faculdade de Farmácia, Universidade de Lisboa, Lisboa, Portugal.
**5** NOVA Institute for Medical Systems Biology, NIMSB, Universidade Nova de Lisboa, 1099-085 Lisboa, Portugal
**6** Instituto Gulbenkian de Ciência, Oeiras, Portugal.
**7** Instituto de Biologia Experimental e Tecnológica, Oeiras, Portugal.

### Supplementary Materials and Methods

#### Determination of proximate composition

The proximate composition of standard and berry-enriched diets was determined based on the Association of Official Analytical Chemists (AOAC) standard analysis methods, as previously described[1]. Water content was determined by drying at 103 ± 2ºC and ash at 550 ± 50ºC. Total nitrogen was measured via the Kjeldahl method, with crude protein calculated by multiplying nitrogen content by 6.25. Crude fat was extracted with petroleum ether using Soxhlet, and crude fibre was analysed using the Weende method. Carbohydrates were estimated by subtracting water, ash, protein, fat, and fibre from the total.

#### Extraction of (poly)phenols in the chow

One hundred and fifty milligrams of freeze-dried food samples were extracted with 5 mL of a mixture of methanol:dimethyl sulfoxide:water (40:40:20) plus 0.1% HCl. The samples were sonicated in an ultrasonic bath for 20 min. After extraction, all samples were centrifuged at 4,000 × g for 10 min at RT, and the resulting supernatant was collected and filtered through a 0.45 µm polyvinylidene difluoride (PVDF) filter before analysis by High-Performance Liquid Chromatography- mass spectrometry (HPLC-MS/MS). Three replicates were extracted and analysed for each diet (normal or rich in berries).

#### HPLC-ESI-IT-MS/MS analysis of the phenolic compounds in diets

HPLC analyses were performed on an Agilent 1200 HPLC system with Electrospray Ionisation-Ion Trap (ESI-IT) mass spectrometer detector in series (Bruker Daltonik, Bremen, Germany). A reverse-phase Pursuit XRs C18 column (250 × 4.0 mm, 5 µm) (Agilent Technologies) was used for the chromatographic separation with water:formic acid (99:1) (phase A) and acetonitrile (phase B) as mobile phases at 0.8 mL/min with the following elution profile: 0 min 5% phase B; 0–15 min, 5–20% phase B; 15–30 min, 20–55% phase B; and 30–45 min 55–90% phase B. Finally, phase B content was decreased to the initial conditions (5%) in 3 min, and the column was re-equilibrated for 6 min. The sample injection volume was 20 µL. The UV-Vis spectra were acquired in the range of 200 to 600 nm. In the mass spectrometer, nitrogen was used as drying and nebulising gas with a pressure of 65 psi, a flow of 11 L/min, and a temperature of 350ºC. The capillary voltage was 4000 V. MS spectra were recorded in negative and positive mode (for the analysis of anthocyanins) with a full scan acquisition in the m/z range 100–1100 with a target mass of 500 m/z. The auto-MS mode was applied to obtain information about the fragmentation patterns.

The compounds were identified by their elution order, UV spectra, molecular weight, fragmentation by MS/MS, and, whenever possible, chromatographic comparison with authentic standards. External calibration curves with appropriate standards belonging to the different families of phenolic compounds were used for the quantification. Cyanidin-3-*O*-glucoside chloride (at 520 nm) was used for anthocyanins quantification, ellagic acid (at 360 nm) for ellagic acid derivatives, catechin (at 280 nm) for flavan-3-ols, chlorogenic acid (at 320 nm) for hydroxycinnamic acids at 320 nm and rutin at 360 nm for flavonols.

**Interference test**

Keeping in mind the complex toxicokinetics of the MPTP, we performed an interference test to ensure that the observed effects of berry supplementation were related to a genuine and meaningful molecular event and not to interference with the pharmacology of the toxin. Since the peak of the 1-methyl-4-phenylpyridinium ($MPP^+$), the toxic metabolite of the neurotoxin MPTP, has been described to be found approximately 90 min after acute MPTP administration[2], a total of $n$=6-8 animals from the MPTP and B+ MPTP groups were sacrificed 90 min after the last MPTP injection, and the blood and striatum were collected. The concentration of MPTP in the blood and $MPP^+$ in the striatum was quantified using HPLC with a UV detector, as previously described[2].

**Determination of Total Phenolic Content**

The Total Phenolic Content (TPC) was quantified using the Folin-Ciocalteu's method as previously described[3]. (Poly)phenols from foods were extracted by mixing 100 mg of the sample with 1 mL of 95% methanol, followed by centrifugation at 10,000 × g for 10 min at room temperature (RT). The supernatant was collected for phenolic content analysis. Briefly, 5 µL of the sample, 15 µL Folin-Ciocalteu's reagent, and 235 µL water were prepared in a 96-well plate, followed by the addition of 45 µL of 20% $Na_2CO_3$ at 40ºC. After 30 min of dark incubation at RT, the absorbance was recorded at 765 nm. TPC was determined using a gallic acid standard curve (0–700 µg/mL) and expressed in mg of gallic acid equivalents (GAE) per 100 g dry sample.

**Fluoro-Jade C staining**

Neurodegeneration in the brain was analysed using Biosensis ® Ready-to-Dilute™ Fluoro-Jade® C (FJC) Staining Kit (Cat No: TR-100-FJT) following the protocol provided by the manufacturer. Imaging was done using a 10x objective in a Zeiss AxioObserver microscope. The number of FJC-positive cells was manually counted in the SNpc of each section. The data are presented as an average number of degenerating cells per section in the SNpc of each experimental group. A blinded observer analysed FJC immunostaining.

**TUNEL staining**

The apoptosis levels were obtained using a In Situ Cell Death Detection Kit, TMR red (Roche, 12 156 792 910). The terminal deoxynucleotidyl transferase dUTP nick end (TUNEL) reaction preferentially labels DNA strand breaks generated during apoptosis. This allows discrimination of apoptosis from necrosis. The TUNEL assay was done as recommended by the producer. The brain slices were incubated for 30 min at 37ºC with 3 U/mL DNase I recombinant, grade I, dissolved in 10 mM Tris-HCl, pH 7.4, containing 10 mM NaCl, 5 mM $MnCl_2$, 0.1 mM $CaCl_2$, and 25 mM KCl to obtain the positive control for the TUNEL staining. Imaging was done using a 10x objective in a Zeiss AxioObserver microscope.

**TH/MAP2 and Iba1/TMEM119 immunostaining**

Before immunoreaction, the sections were rinsed in 0.01 M PBS for 10 min. The antigen retrieval step was done by immersing the slides for 30 min in citrate buffer (pH 6.0) heated to 75ºC. Three washes were performed with 0.01 M PBS between each of the two consecutive steps during the process. Sections were treated with 1% BSA in 0.01 M PBS in a humid condition for 1h at RT to block nonspecific immunostaining. Subsequently, sections were immunolabelled with mouse anti-TH (Tyrosine hydroxylase) antibody (1:250, #2928, Sigma Aldrich) or chicken anti-MAP2 (Microtubule associated protein-2) antibody (1:1000, #Ab5392, Abcam) in 1% BSA overnight at 4ºC. Following three washes in 0.01 M PBS, sections were incubated with the appropriate secondary antibodies (Alexa Fluor Goat anti-mouse 555; # A-21422 and Alexa Fluor Goat anti-chicken; #11039, Invitrogen) in dilution 1:1000 in 1xPBS 1h in room temperature. DAPI (4′,6-diamidino-2-phenylindole) stain (1:10000, #D9542, Sigma, Aldrich) was used to label the cell nuclei. Following three washes in 0.01 M PBS, coverslips were placed on the sections using Vectashield anti-fading fluorescent mounting media (#H-1000-10, Vector Laboratories). Images were acquired using a Zeiss LSM 710 confocal point-scanning microscope (Zeiss, Germany).

**Quantification of the TH/MAP2-positive cell ratio in the SNpc**

Analysis was done using Z-stack images. The number of TH- and MAP2-positive cells was counted throughout the SNpc region and expressed as the number of TH-positive cells divided by the number of MAP/2/positive cells per section. The number presented as a final result represents an average number of TH-positive/MAP-2-positive cells per section. The blinded observer performed the analysis of the images obtained for all histological and immunolabeling.

**Iba1 and GFAP immunostaining**

Before immunoreaction, the sections were rinsed in 0.01 M PBS for 10 min and treated with 0.3% hydrogen peroxide in 20% methanol solution for 30 min at RT to block endogenous peroxidase. Three washes were performed with 0.01 M PBS between the two consecutive steps during the process. Sections were treated with 3% BSA in 0.01 M PBS in a humid condition for 1h at RT to block nonspecific immunostaining. Subsequently, sections were immunolabelled with rabbit anti-

Iba1 (ionized calcium-binding adaptor molecule-1) antibody (1:500, Cat.No.019-19741, Wako, Japan) or rabbit anti-GFAP (Glial fibrilar acidic protein) antibody (1:2000, Cat.No. Z0334, Dako) in 0.1% Triton X-100/0.01 M PBS overnight at 4ºC. Following three washes in 0.01 M PBS, sections were incubated with the EnVision+ System-HRP Labelled Polymer Anti-Rabbit (K4003, DAKO, Denmark) in 0.01 M PBS in a humid condition for 2h at RT. Bound antibodies were visualised with DAB (3,3-diaminobenzidine tetrahydrochloride; Dako, Agilent Technologies, Denmark) by the avidin-biotin-peroxidase complex method, following standard protocols (Vector Laboratories, Burlingame, CA). All sections were dehydrated in graded ethanol, cleared in xylene, and mounted in Canada balsam (Merck, USA). A negative control slide was treated the same way but with the omission of the primary antibody and run alongside the other samples to test the specificity of the reaction. Imaging was done using a 5x, 10x or 40x objective in a Zeiss AxioObserver microscope.

**Quantification of the number and morphology of Iba1-positive cells**

For counting Iba1-positive cells, three rectangles (500 µm x 1000 µm) were superimposed over the image for each brain section, covering different parts of the area of interest. The number of Iba1-positive cells was counted for each region of interest (ROI), and the average number of cells was calculated per section.

For the Iba1-positive cells morphology, cell bodies were manually outlined regarding Iba1-positive cell soma size, and their area was determined using NIH ImageJ 1.50i (US National Institutes of Health, Bethesda, USA). A total of $n \geq 100$ cells per animal/per specific region ($n \geq 400$ cells per experimental group/per specific region) was analysed.

**Quantification of the GFAP staining intensity**

Images obtained using a 5x objective in a Zeiss AxioObserver microscope were converted to 8-bit images, and the ROIs (motor cortex, striatum and SN) were manually outlined and analysed using the Uncalibrated optical density option in the ImageJ 1.50i program. For the SN, the same ROI was superposed to all images circling the entire SN region and mean grey intensity levels were obtained. Mean grey intensity levels were obtained.

### Supplementary Results

**Table S1**. Proximate composition of diets based on the standard methods of the Association of Official Analytical Chemists (AOAC).

| | | Standard diet | Berry-enriched diet |
|---|---|---|---|
| **Energy** | kJ / g | 15.81 ± 0.17 | 15.35 ± 0.11 |
| | kcal / g | 3.76 ± 0.04 | 3.65 ± 0.03 |
| **Water** | % | 5.14 ± 0.09 | 7.52 ± 0.05 |
| **Dry matter** | | 94.86 ± 0.09 | 92.48 ± 0.04 |
| **Carbohydrates** | g/100g | 63.52 ± 0.58 | 62.27 ± 0.36 |
| **Proteins** | | 18.47 ± 0.20 | 17.07 ± 0.06 |
| **Lipids** | | 4.70 ± 0.02 | 4.51 ± 0.01 |
| **Fibres** | | 3.07 ± 0.37 | 3.74 ± 0.39 |
| **Ashes** | | 5.09 ± 0.06 | 4.89 ± 0.07 |

**Table S2.** Phenolic compounds quantified by HPLC-UV-IT in the berry-supplemented diet.

| Compounds | RT | $m/z^+$ | MS/MS | $\lambda_{max}$ | mg/100 g |
|---|---|---|---|---|---|
| **Total *Anthocyanins:*** | | | | | **30.85 ± 3.59** |
| Delphinidin-*O*-galactoside | 12.10 | 465 | 303 | 524 | 1.22 ± 0.08 |
| Delphinidin-*O*-glucoside | 13.30 | 465 | 303 | 524 | 5.32 ± 0.93 |
| Cyanidin-*O*-galactoside | 14.00 | 449 | 285 | 522 | 7.18 ± 1.01 |
| Cyanidin-3-*O*-glucoside* | 15.20 | 449 | 285 | 518 | 10.58 ± 0.92 |
| Petunidin-*O*-glucoside | 16.00 | 479 | 317 | 526 | 1.12 ± 0.04 |
| Malvidin-*O*-glucoside | 17.40 | 493 | 331/315 | 520 | 3.57 ± 0.51 |
| Peonidin-*O*-glucoside | 18.90 | 463 | 301 | 526 | 1.86 ± 0.10 |
| | | $m/z^-$ | | | |
| ***Total Hydroxycinnamic acids:*** | | | | | **6.72 ± 0.12** |
| Chlorogenic acid | 14.60 | 353 | 191 | 296/312 | 3.05 ± 0.02 |
| Di-*O*-caffeoylquinic acid | 19.70 | 515 | 353/335 | 298/326 | 2.10 ± 0.07 |
| Caffeic acid | 23.90 | 179 | - | 296/322 | 1.57 ± 0.03 |
| ***Flavon-3-ols*** | | | | | |
| Catechin* | 17.3 | 289 | 245 | 280 | **8.69 ± 1.54** |
| ***Ellagic acid derivatives*** | | | | | |
| Ellagic acid | 22.90 | 301 | 256/184 | 254/360 | **11.92 ± 2.56** |
| ***Flavonols*** | | | | | **3.14 ± 0.66** |
| Myricetin-*O*-glucoside | 20.10 | 479 | 316 | 258/362 | 0.35 ± 0.01 |
| Quercetin-3-*O*-rutinoside (rutin)* | 22.00 | 609 | 301 | 254/350 | 1.29 ± 0.31 |
| Quercetin-*O*-arabinoside | 24.70 | 433 | 301 | 248/348 | 0.66 ± 0.19 |
| Quercetin-*O*-rhamnoside | 25.70 | 447 | 301 | 248/350 | 0.84 ± 0.15 |

*Quantified with their authentic standards. The estimated limits of detection (LOD) and quantification (LOQ) in ppm for the selected phenolic compounds were as follows: for cyanidin 3-glucoside, LOD = 7.42 ppm and LOQ = 22.47 ppm; for catechin, LOD = 9.58 ppm and LOQ = 29.03 ppm; for ellagic acid, LOD = 9.97 ppm and LOQ = 30.22 ppm; and for Quercetin 3-O-rutinoside (rutin) LOD = 20.15 ppm and LOQ = 61.05 ppm.

**Table S3.** Phenolic compounds quantified by HPLC-UV-IT in the standard diet.

| Compounds | RT | *m/z*$^-$ | MS/MS | $\lambda_{max}$ | mg/100 g |
|---|---|---|---|---|---|
| ***Total Hydroxycinnamic acids:*** | | | | | **3.3 ± 0.1** |
| Di-*O*-caffeoylquinic acid | 19.70 | 515 | 353/335 | 298/326 | 2.75 ± 0.09 |
| Caffeic acid | 23.90 | 179 | - | 296/322 | 0.55 ± 0.1 |

**Table S4**. List of low-molecular weight polyphenol metabolite names according to recommendations (Curti C et al., 2025) and common names in brackets.

| **Benzoic acids** |
|---|
| 3,4,5-Trihydroxybenzoic acid (Gallic acid) |
| 3,4,5-Trihydroxybenzoic acid-glucuronide isomers (Gallic acid glucuronide isomers) |
| 3,4,5-Trihydroxybenzoic acid-sulfate isomers (Gallic acid sulfate isomers) |
| 3,4-Dihydroxy-5-methoxybenzoic acid (5-Methylgallic acid) |
| Dihydroxy-methoxybenzoic acid-glucuronide isomers (Methylgallic acid glucuronide isomers) |
| Dihydroxy-methoxybenzoic acid-sulfate (Methylgallic acid sulfate) |
| 4-Hydroxy-3,5-dimethoxybenzoic acid (Syringic acid) |
| 3,5-Dimethoxybenzoic acid-4-glucuronide isomers (Syringic acid glucuronide) |
| 3,5-Dimethoxybenzoic acid-4-sulfate isomers (Syringic acid sulfate) |
| Benzoic acid |
| Benzoic acid-glucuronide isomers |
| Benzoic acid-sulfate isomers |
| Hydroxybenzoic acid isomers |
| Hydroxybenzoic acid-glucuronide isomers |
| Hydroxybenzoic acid-sulfate isomers |
| Dihydroxybenzoic acid isomers |
| Dihydroxybenzoic acid-glucuronide isomers |
| Dihydroxybenzoic acid-sulfate isomers |
| 2,3-Dimethoxybenzoic acid (Veratric acid) |
| Hydroxy-methoxybenzoic acid isomers (vanillic/ isovanillic acid isomers) |
| Methoxybenzoic acid-glucuronide isomers (vanillic/isovanillic acid-glucuronide isomers) |
| Methoxybenzoic acid-sulfate isomers (vanillic/isovanillic acid-sulfate isomers) |
| 3-(4′-Hydroxyphenoxy)benzoic acid |
| 3-(4’-Hydroxyphenoxy)benzoic acid glucuronide |
| 3-(4’-Hydroxyphenoxy)benzoic acid sulfate |
| Hydroxybenzoic acid isomers |
| Hydroxybenzoic acid glucuronide isomers |
| Hydroxybenzoic acid sulfate isomers |
| **Phenylacetic acids, Mandelic acids, 3-(phenyl)propanoic acids, and cinnamic acids** |
| Phenylacetic acid |
| Phenylacetic acid-glucuronide isomers |
| Acetylphenol-sulfate isomers |
| 2-(Hydroxy-methoxyphenyl)acetic acid isomers (homovanillic acid/homoisovanillic acid) |
| 2-(Methoxyphenyl)acetic acid-glucuronide isomers (homovanillic/homoisovanillic acid glucuronide isomers) |
| 2-(Methoxyphenyl)acetic acid-sulfate isomers (homovanillic/homoisovanillic acid sulfate isomers) |

3′,4′-Dihydroxyphenylacetic acid (Homoprotocatechuic acid or DOPAC)

3′,4′-Dihydroxyphenylacetic acid-glucuronide isomers (Homoprotocatechuic acid glucuronide isomers)

3′,4′-Dihydroxyphenylacetic acid-sulfate isomers (Homoprotocatechuic acid acid sulfate isomers)

2-Hydroxy-2-(phenyl)acetic acid (Mandelic acid)

2-Hydroxy-2-(phenyl)acetic acid-glucuronide isomers (Mandelic acid glucuronide isomers)

2-Hydroxy-2-(phenyl) acetic acid-sulfate isomers (Mandelic acid sulfate isomers)

2-Hydroxy-2-(4′-hydroxy-3′-methoxyphenyl)acetic acid (vanililmandelic acid)

2-Hydroxy-2-(hydroxy-methoxyphenyl)acetic acid-glucuronide isomers (vanililmandelic acid glucuronide isomers)

2-Hydroxy-2-(hydroxymethoxyphenyl)acetic acid-sulfate isomers (vanililmandelic acid sulfate isomers)

3′,4′-Dihydroxycinnamic acid (Caffeic acid)

3′,4′-Dihydroxycinnamic acid acid-glucuronide isomers (Caffeic acid glucuronide isomers)

3′,4′-Dihydroxycinnamic acid acid-sulfate isomers (Caffeic acid sulfate isomers)

Hydroxy-methoxycinnamic acid (Ferulic/isoferulic acid isomers)

Methoxycinnamic acid-glucuronide (Ferulic/isoferulic acid glucuronide isomers)

Methoxycinnamic acid-sulfate (Ferulic/isoferulic acid sulfate isomers)

3-(3′,4′-Dihydroxyphenyl)propanoic acid (Dihydrocaffeic acid)

3-(3′,4′-Dihydroxyphenyl)propanoic acid-glucuronide isomers (Dihydrocaffeic acid glucuronide isomers)

3-(3′,4′-Dihydroxyphenyl)propanoic acid-sulfate isomers (Dihydrocaffeic acid sulfate isomers)

3-(Hydroxyphenyl)propanoic acid isomers

3-(Hydroxyphenyl)propanoic acid acid-glucuronide (Hydroxy-hydrocinnamic acid glucuronide isomers)

3-(Hydroxyphenyl)propanoic acid-sulfate (Hydroxy-hydrocinnamic acid sulfate isomers)

3-(4′-Hydroxy-3′-methoxyphenyl)propanoic acid (Dihydroferulic acid)

3-(4′-Hydroxy-3′-methoxyphenyl)propanoic acid-glucuronide (Dihydroferulic acid glucuronide)

3-(4′-Hydroxy-3′-methoxyphenyl)propanoic acid-sulfate (Dihydroferulic acid sulfate)

3-(3′-Hydroxy-4′-methoxyphenyl)propanoic acid (Dihydroisoferulic acid)

3-(3′-Hydroxy-4′-methoxyphenyl)propanoic acid-sulfate (Dihydroisoferulic acid sulfate)

3-(Phenyl)propanoic acid (Hydrocinnamic acid)

3-(Phenyl)propanoic acid-sulfate (Hydrocinnamic acid sulfate isomers)

**Hippuric acids**

Hippuric acid (Hippuric acid)

Hydroxyhippuric acid isomers

Hydroxyhippuric acid-glucuronide isomers

Hydroxyhippuric acid-sulfate isomers

**Benzaldehydes**

| |
|---|
| Hydroxybenzaldehyde isomers |
| Benzaldehyde-glucuronide isomers |
| Benzaldehyde-sulfate isomers |
| Dihydroxybenzaldehyde isomers |
| Dihydroxybenzaldehyde-glucuronide isomers |
| Dihydroxybenzaldehyde-sulfate isomers |
| 2,4,6-trihydroxybenzaldehyde (phloroglucinaldehyde) |
| Trihydroxybenzaldehyde-glucuronide isomers |
| Trihydroxybenzaldehyde-sulfate isomers |
| Hydroxy-methoxybenzaldehyde (Vanillin/ isovanillin) isomers |
| Hydroxy-methoxybenzaldehyde-glucuronide isomers (Vanillin/ isovanillin glucuronide isomers) |
| Hydroxy-methoxybenzaldehyde-sulfate isomers (Vanillin/ isovanillin sulfate isomers) |
| **Benzene diols and triols** |
| Benzene-1,2-diol (Catechol) |
| 2-hydroxyphenyl hydrogen glucuronide (Catechol glucuronide) |
| 2-hydroxyphenyl hydrogen sulfate (Catechol sulfate) |
| 2-methoxyphenol (Methylcatechol or guaiacol) |
| 2-methoxyphenol-glucuronide (guaiacol glucuronide) |
| 2-methoxyphenol-sulfate (guaiacol sulfate) |
| Benzene-1,3-diol (Resorcinol) |
| 3-hydroxyphenyl hydrogen glucuronide isomers (Resorcinol glucuronide) |
| 3-hydroxyphenyl hydrogen sulfate isomers (Resorcinol sulfate) |
| Benzene-1,2,3-triol (Pyrogallol) |
| Dihydroxybenzene-glucuronide isomers (Pyrogallol glucuronide isomers) |
| Dihydroxybenzene-sulfate isomers (Pyrogallol sulfate isomers) |
| Benzene-1,3,6-triol (Phloroglucinol) |
| Dihydroxybenzene-glucuronide isomers (Phloroglucinol glucuronide isomers) |
| Dihydroxybenzene-sulfate isomers (Phloroglucinol sulfate isomers) |
| Methoxy-dihydroxybenzene (methylpyrogallol or methoxypyrogallol) |
| Methoxy-hydroxybenzene-glucuronide isomers (Methylpyrogallol glucuronide isomers) |
| Methoxy-hydroxybenzene-sulfate isomers (Methylpyrogallol sulfate isomers) |
| **Flavan-3-ols, 5-(phenyl)-gamma-valerolactones, and 5-(phenyl)valeric acids** |
| Epicatechin |
| Epicatechin-glucuronide isomers |
| Epicatechin-sulfate isomers |
| Epicatechin-sulfate-glucuronide |
| Methyl-*O*-Epicatechin |
| Methyl-*O*-epicatechin-glucuronide isomers |
| Methyl-*O*-epicatechin-sulfate isomers |

| |
|---|
| 3′,4′-Dimethyl-*O*-epicatechin |
| 3′,4′-Dimethyl-*O*-epicatechin-glucuronide isomers |
| 3′,4′-Dimethyl-*O*-epicatechin-sulfate isomers |
| 5-(Phenyl)-γ-valerolactone |
| 5-(3′,4′-Dihydroxyphenyl)-γ-valerolactone isomers |
| 5-(3′,4′-Dihydroxyphenyl)-γ-valerolactone glucuronide isomers |
| 5-(3′,4′-Dihydroxyphenyl)-γ-valerolactone sulfate isomers |
| 5-(Hydroxyphenyl)-γ-valerolactone isomers |
| 5-(Hydroxyphenyl)-γ-valerolactone glucuronide isomers |
| 5-(Hydroxyphenyl)-γ-valerolactone sulfate isomers |
| 5-(Phenyl)-γ-valerolactone glucuronide isomers |
| 5-(Phenyl)-γ-valerolactone sulfate isomers |
| 5-(Hydroxyphenyl)valeric acid isomers |
| 5-(3′,4′-Dihydroxyphenyl)valeric acid isomers |
| 5-(3′,4′-Dihydroxyphenyl)valeric acid-glucuronide isomers |
| 5-(3′,4′-Dihydroxyphenyl)valeric acid-sulfate isomers |
| 5-(Hydroxyphenyl)valeric acid-glucuronide isomers |
| 5-(Hydroxyphenyl)valeric acid-sulfate isomers |
| 5-(Phenyl)valeric acid |
| 5-(Phenyl)valeric acid-glucuronide isomers |
| 5-(Phenyl)valeric acid-sulfate isomers |
| **1-(3′,4′-dihydroxyphenyl)-3-(2″,4″,6″-trihydroxyphenyl)-2-propanol (3,4-diHHP-2-ol)** |
| 1-(3′,4′-dihydroxyphenyl)-3-(2″,4″,6″-trihydroxyphenyl)-2-propanol (3,4-(dihydroxyphenyl)-2-propanol) |
| 1-(3′,4′-dihydroxyphenyl)-3-(2″,4″,6″-trihydroxyphenyl)-2-propanol- glucuronide isomers (3,4-(dihydroxyphenyl)-2-propanol glucuronide isomers) |
| 1-(3′,4′-dihydroxyphenyl)-3-(2″,4″,6″-trihydroxyphenyl)-2-propanol- sulafte isomers (3,4-(dihydroxyphenyl)-2-propanol-2-ol-sulfate isomers) |
| 1-(hydroxyphenyl)-3-(2″,4″,6″-trihydroxyphenyl)-2-propanol isomers (3/4-(hydroxyphenyl)-2-propanol isomers) |
| 1-(phenyl)-3-(2″,4″,6″-trihydroxyphenyl)-2-propanol- glucuronide isomers (3-(hydroxyphenyl)-2-propanol-glucuronide isomers) |
| 1-(phenyl)-3-(2″,4″,6″-trihydroxyphenyl)-2-propanol- sulfate isomers (3-(hydroxyphenyl)-2-propanol-sulfate isomers) |

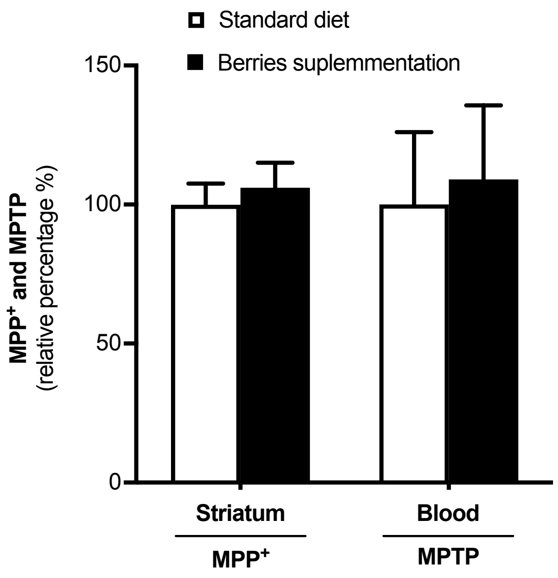


**Figure S1.** MPTP and $MPP^+$ quantification in blood and striatal tissue. Mice (*n*=5-8 per group) were fed for 6 weeks (standard or berry-enriched diet) and treated with 4x15 mg MPTP/kg of body weight. Striatum and blood were collected 90 min after the last injection of MPTP. Samples were processed for MPTP levels in the blood and $MPP^+$ in the striatum, as described in the methods.

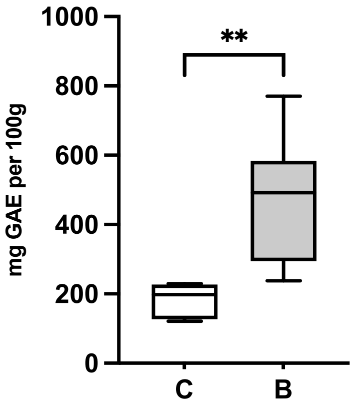


**Figure S2.** Total Phenolic Content (TPC) in the standard diet (C) and berry-supplemented diet (B), which was quantified by using Folin-Ciocalteu's method. TPC was determined using a gallic acid standard curve (0-700 µg/mL) and expressed in mg of gallic acid equivalents (GAE) per 100 g dry sample. Statistical significance: **p<0.01.

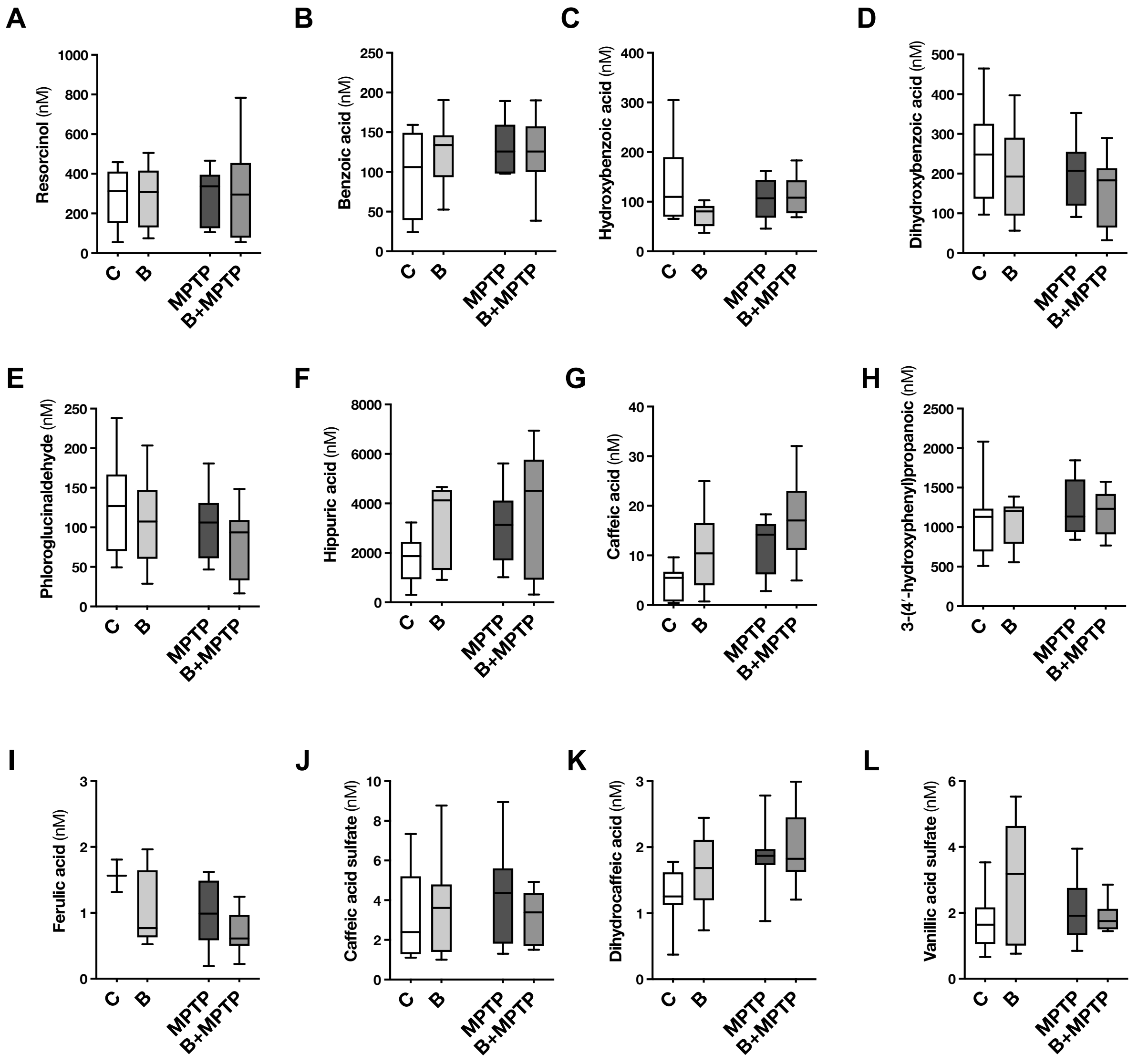


**Figure S3. Quantification of plasma metabolites found in circulation, whose standards were available.** Quantification of (**A**) Resorcinol, (**B**) Benzoic acid, (**C**) Hydroxybenzoic acid, (**D**) Dihydroxybenzoic acid, (**E**) Phloroglucinaldehyde, (**F**) Hippuric acid, (**G**) Caffeic acid, (**H**) 3-(4′-hydroxyphenyl)propanoic, (**I**) Ferulic acid, (**J**) Caffeic acid sulfate, (**K**) Dihydrocaffeic acid, and (**L**) Vanillic acid sulfate. (*n*=8-10). C – Control group; MPTP – MPTP group; B – Berry group; B+MPTP – Berry+MPTP group.

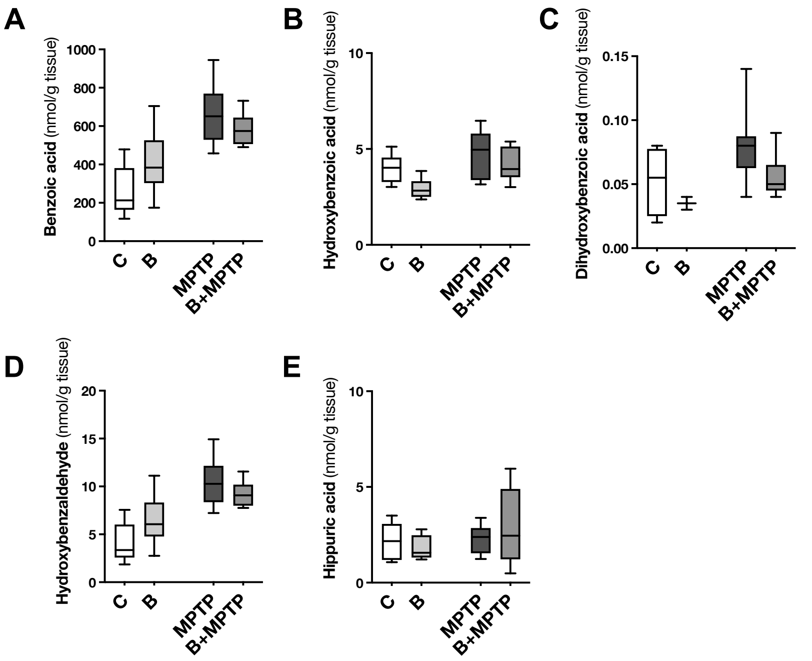


**Figure S4. Quantification of metabolites in perfused brains whose standards were available.** Quantification of (**A**) Benzoic acid, (**B**) Hydroxybenzoic acid, (**C**) Dihydroxybenzoic acid, (**D**) Hydroxybenzaldehyde, and (**E**) Hippuric acid. *n*=8-10. C – Control group; MPTP – MPTP group; B – Berry group; B+MPTP – Berry+MPTP group.

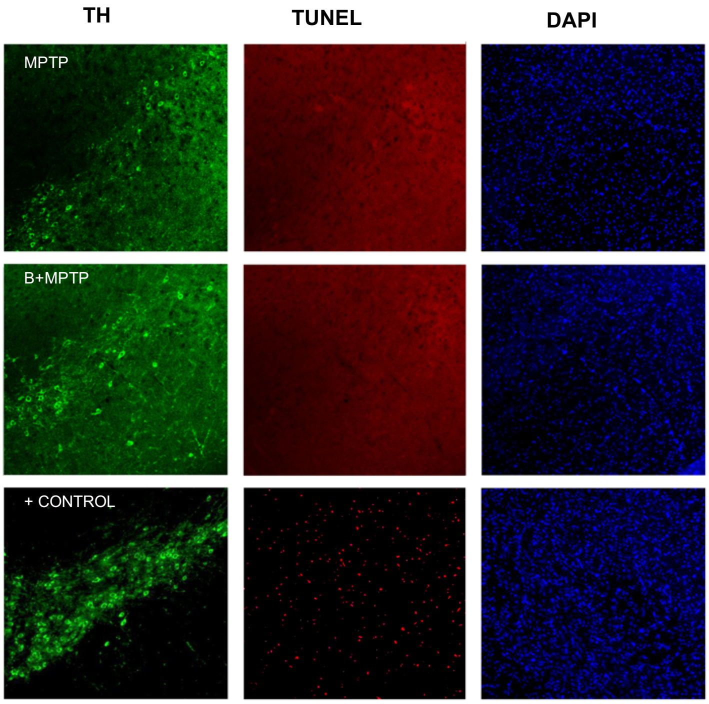


**Figure S5.** TUNEL assay of SN region of MPTP (first row) and B+MPTP (second row) animals 24h post-MPTP injection. Triple immunofluorescent staining of TH-labelled neurons (left column, green), TUNEL assay (middle column, red) and DAPI (right column, blue). No TUNEL-positive nuclei were observed in the SN of either MPTP or B+MPTP animals. Positive control (last row) where the brain tissue was treated with a high concentration of DNAse1 shows numerous TUNEL-positive nuclei (middle image, red). MPTP – MPTP group; B+MPTP – Berry+MPTP group.

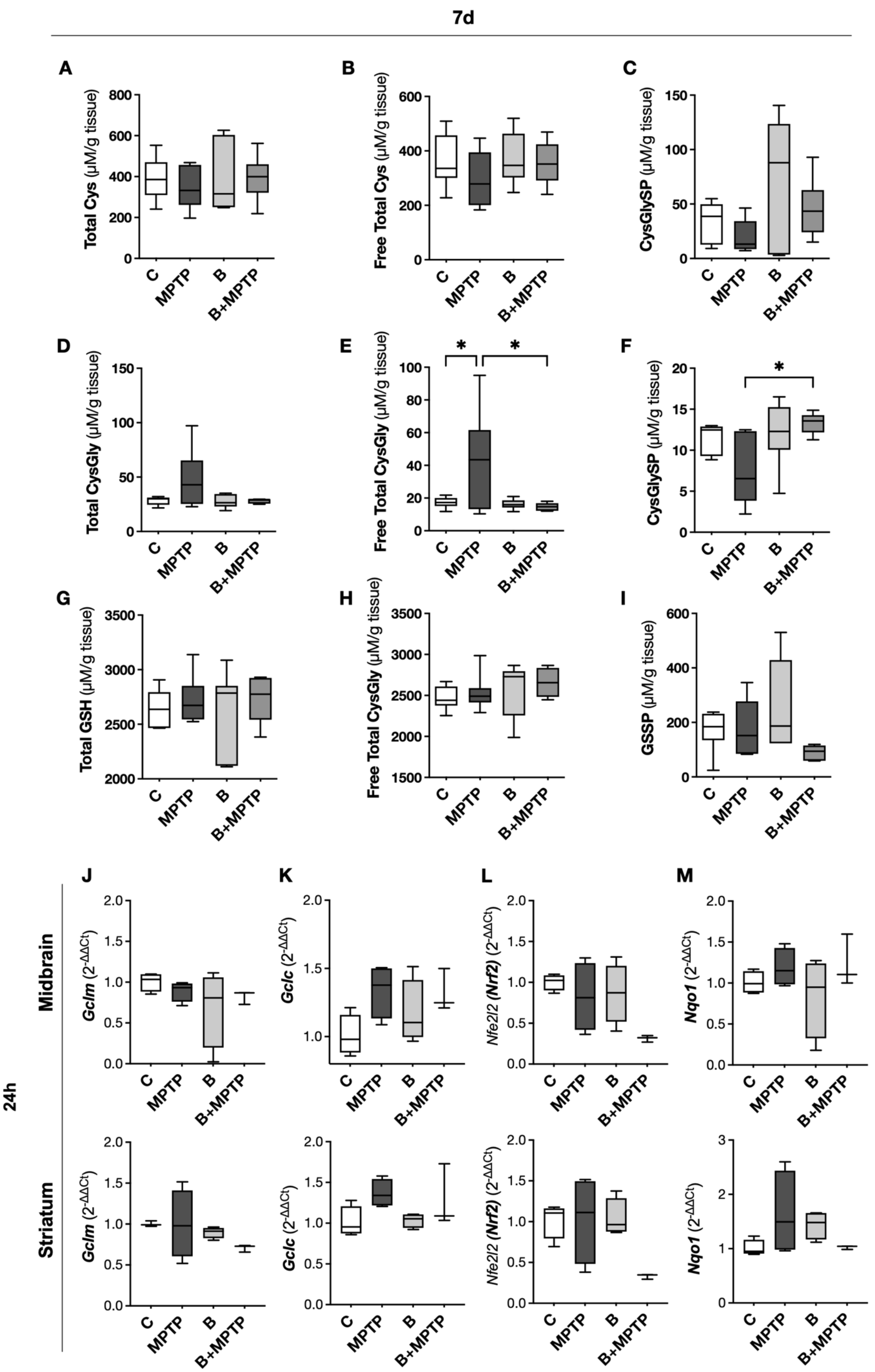
7d
A
Total Cys (µM/g tissue)
B
Free Total Cys (µM/g tissue)
C
CysGlySP (µM/g tissue)
D
Total CysGly (µM/g tissue)
E
Free Total CysGly (µM/g tissue)
F
CysGlySP (µM/g tissue)
G
Total GSH (µM/g tissue)
H
Free Total CysGly (µM/g tissue)
I
GSSP (µM/g tissue)
C
MPTP
B
B+MPTP
*
24h
Midbrain
Striatum
J
K
L
M
Gclm (2^-ΔΔCt)
Gclc (2^-ΔΔCt)
Nfe2l2 (Nrf2) (2^-ΔΔCt)
Nqo1 (2^-ΔΔCt)

**Figure S6.** Cysteine-related thiols in the brain 7d post-MPTP injections. Thiolome profile at 7d: (**A**) Total Cys; (**B**) Free total Cys; (**C**) Cysteinylated proteins (CysSSP); (**D**) Total CysGly; (**E**) Free total CysGly; (**F**) Cysteinylglycinylated proteins (CysGlySP); (**G**) Total GSH; (**H**) Free total GSH; (**I**) Glutathionylated proteins (GSSP). n=5-7 per group. mRNA levels of (**J**) *Gclm*, (**K**) *Gclc*, (**L**) *Nfe2l2* (*Nrf2*), and (**M**) *Nqo1* in the midbrain and striatum at 24h timepoint. n=3-4 per group. Statistical significance: *p<0.05. C – Control group; MPTP – MPTP group; B – Berry group; B+MPTP – Berry+MPTP group.

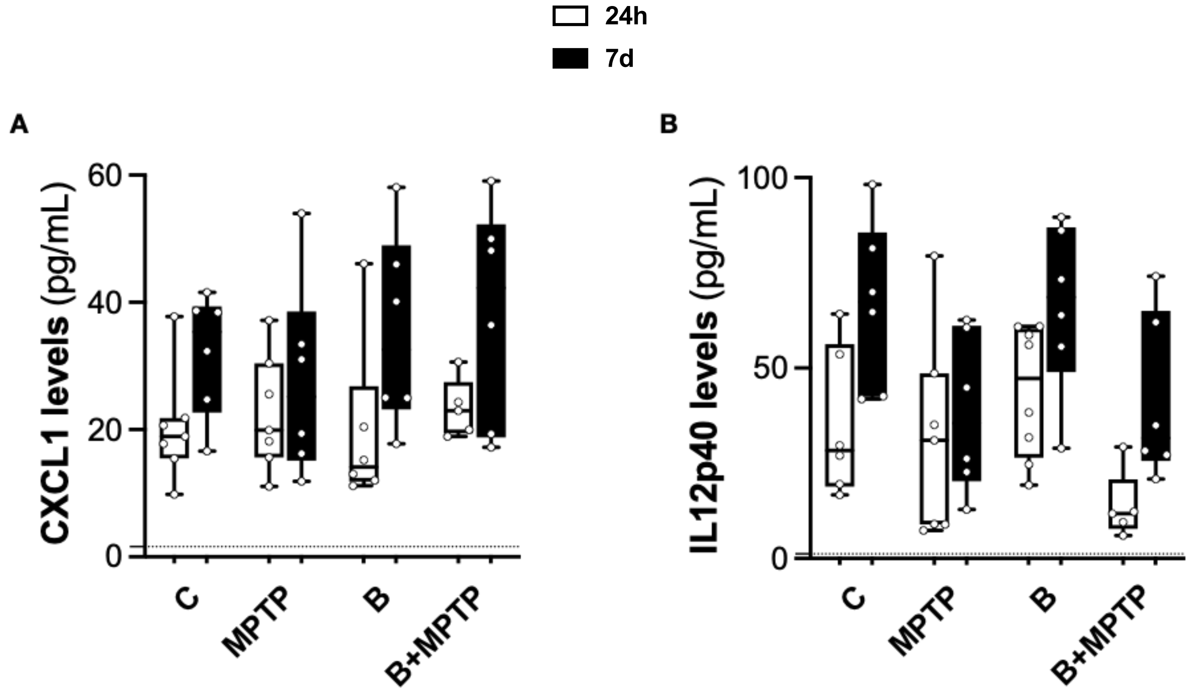


**Figure S7.** Serum levels of (**A**) CXCL1 and (**B**) IL12p40 quantified with the bead-based LEGENDplex immunoassay at 24h and 7d timepoints. The dashed line represents the limit of detection for each molecule. *n*=6-7 per group. C – Control group; MPTP – MPTP group; B – Berry group; B+MPTP – Berry+MPTP group.
