## Supplementary figures and images for "From berries to brain: Assessing the impact of (poly)phenols in the MPTP mouse model of Parkinson’s disease"

### Figure S1

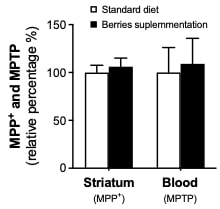

### Figure S2

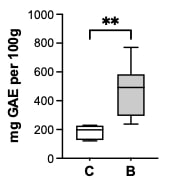

### Figure S3

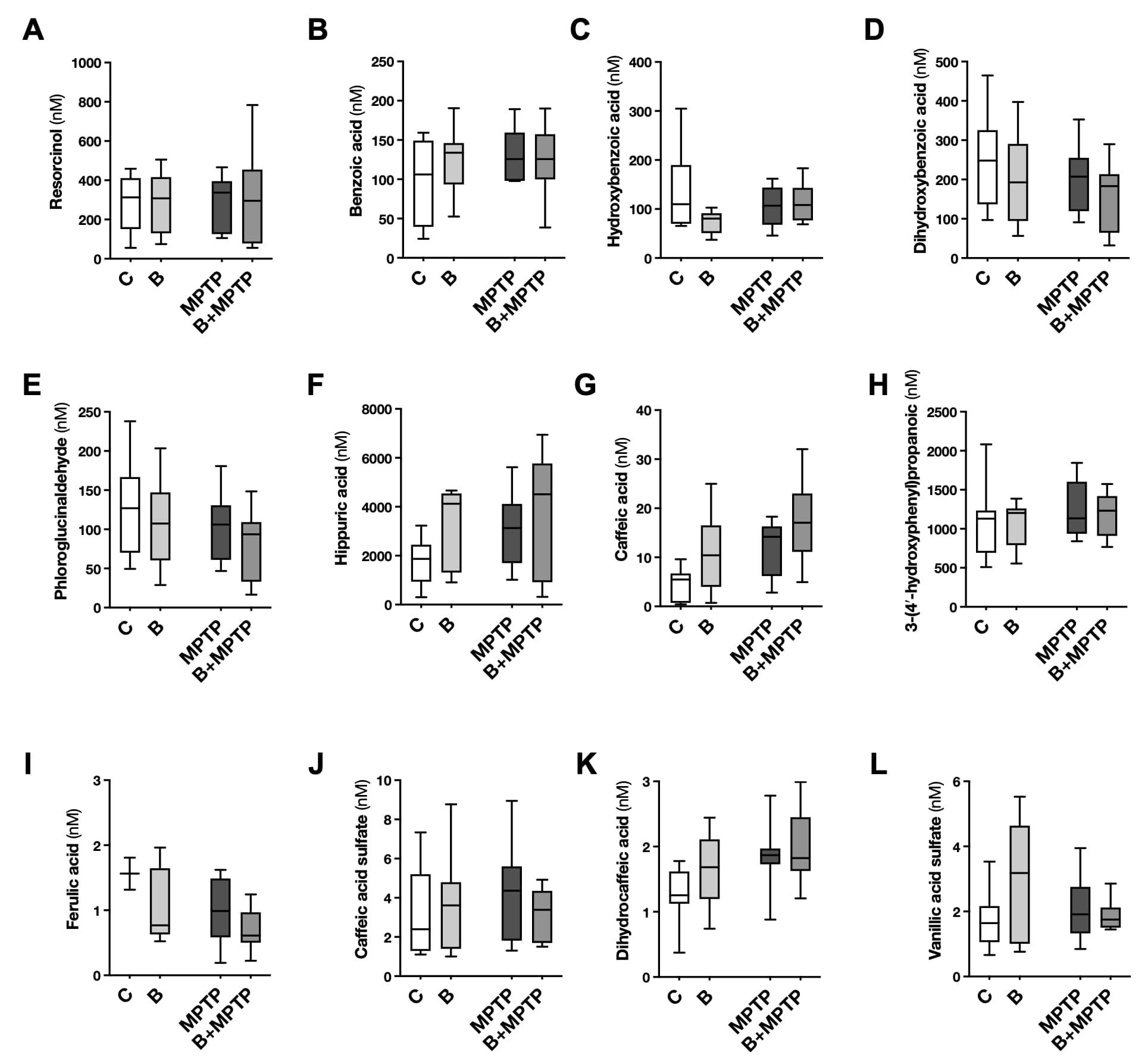

### Figure S4

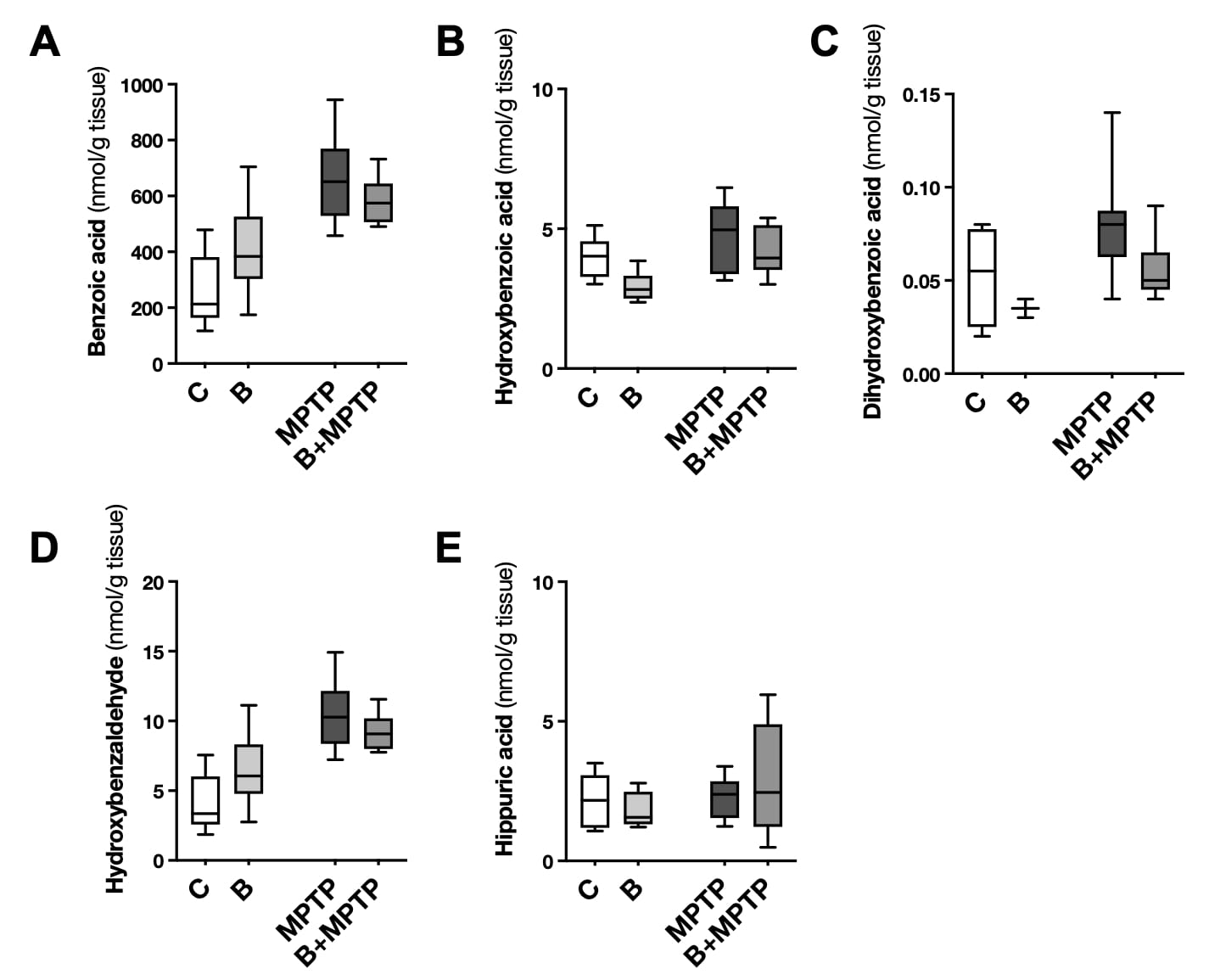

### Figure S5

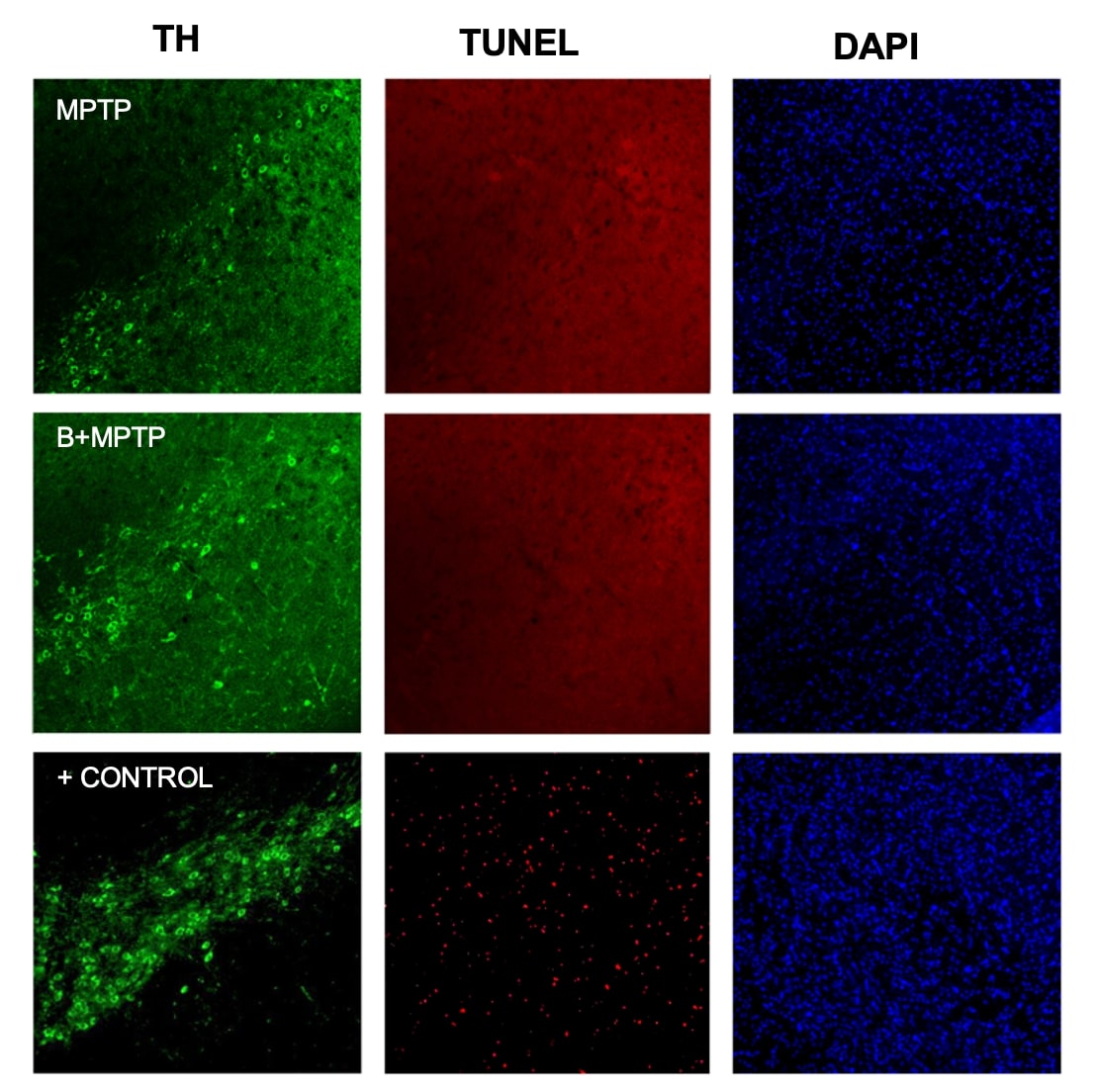

### Figure S6

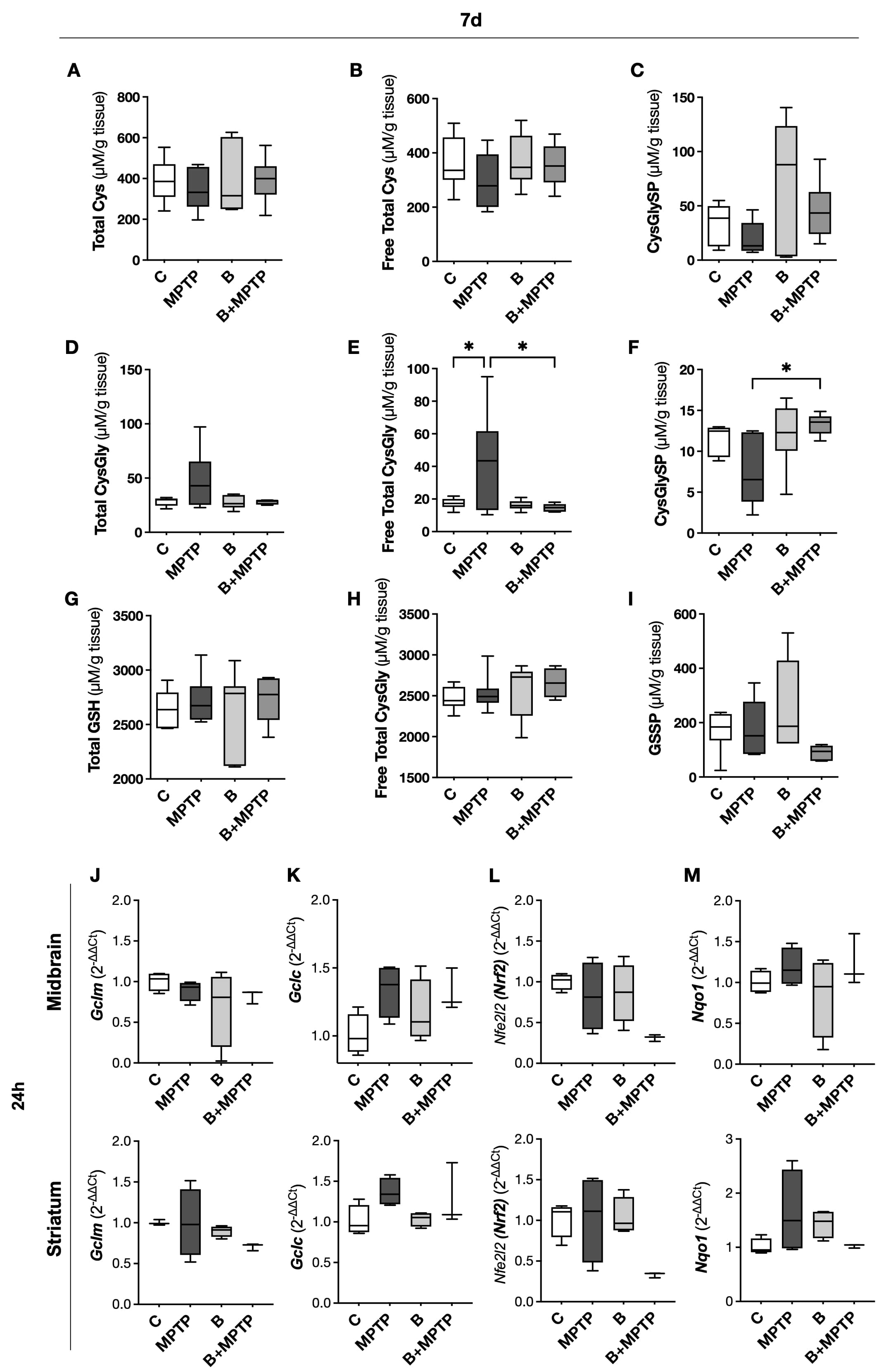

### Figure S7

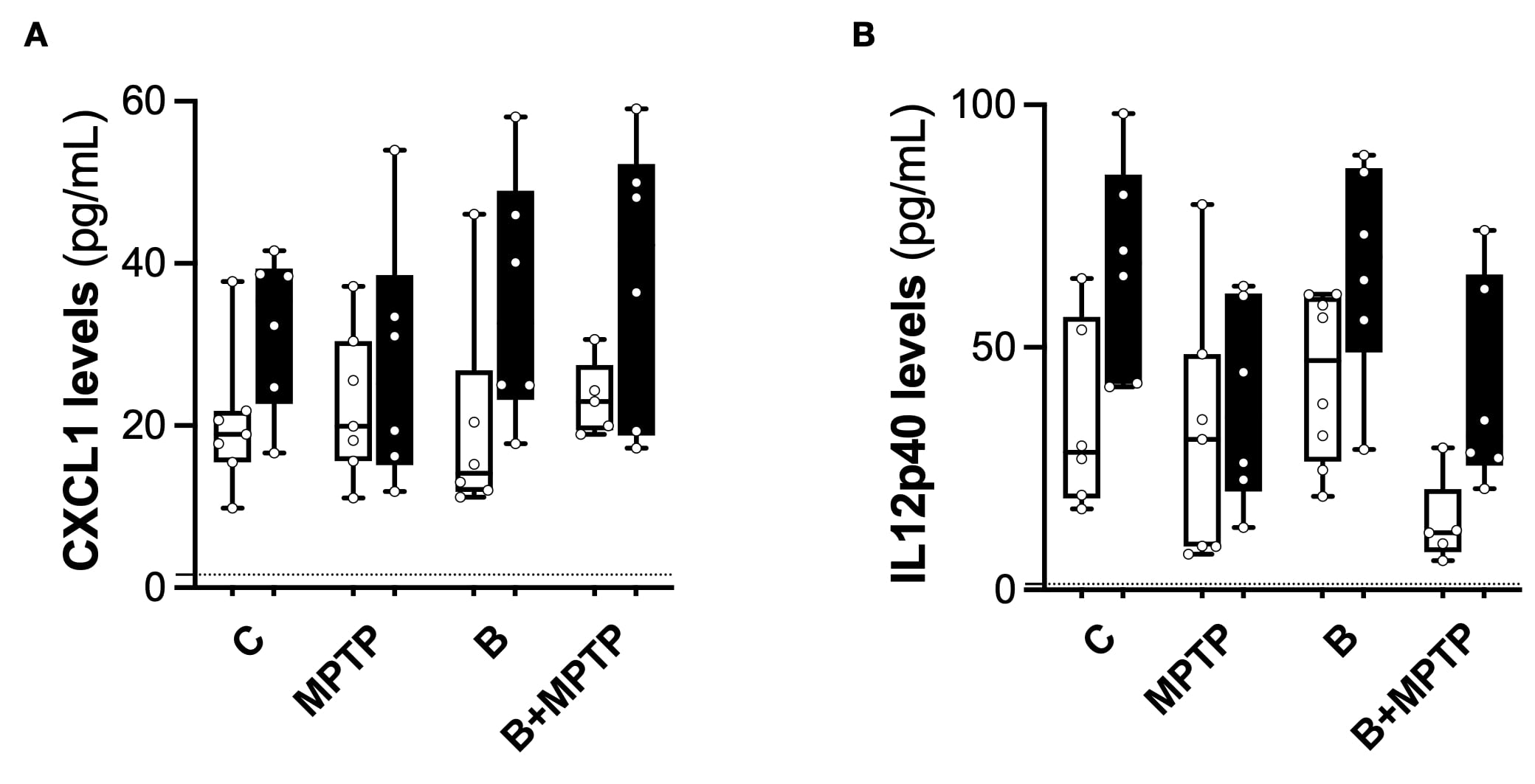
